# Comparative analysis identifies a shared topological organisation among divergent arenavirus intergenic RNAs

**DOI:** 10.64898/2026.10.22.760956

**Authors:** Alex Childs, Dinh Thien Kim Ly, Tycho Marinus, Rahul A Bhosale, Janet M Daly, Katarzyna Tych, Toshana L Foster, Aditi N Borkar

**Affiliations:** One Virology Wolfson Centre for Global Virus Research, School of Veterinary Medicine and Science, University of Nottingham, UK; Groningen Biomolecular Sciences and Biotechnology Institute, University of Groningen, Groningen, the Netherlands; School of Biosciences, University of Nottingham, UK; Institute for Microsystems Engineering (IMTEK), Faculty of Engineering, University of Freiburg, Freiburg im Breisgau, Germany; Institute of Systems, Molecular and Integrative Biology, University of Liverpool, UK

**Keywords:** RNA, topology, intergenic regions, cryo-OrbiSIMS, arenavirus

## Abstract

Biological function is often inferred from sequence conservation or structural similarity. However, many viral RNAs maintain regulatory functions despite extensive diversification of both sequence and structures. Using arenavirus intergenic regions (IGRs) as a model system, we investigated what level of RNA organization remains conserved when these conventional sequence-structure-function relationships break down. Combining cryogenic OrbiSIMS guided RNA structure determination with graph theory-based analysis of fifty-eight IGRs across phylogenetically diverse arenaviruses, we discovered that IGR topology, defined by the connectivity and relative integration of individual stem-loop elements, is consistently arranged around a highly connected helical core that persists across divergent IGR folds. Quantitative centrality analyses further revealed that topologically peripheral stem-loops are comparatively sparse and variable across the phylogenetic classes. Investigation of the functional consequence of the peripheral elements in Golden Gate virus (a reptile arenavirus) S-IGR revealed selective association with interferon-induced and antiviral RNA-binding proteins in human host cells infected with recombinant Lymphocytic Choriomeningitis virus (a distantly related human arenavirus). Taken together, these findings identify graph-defined topology as a recurrent organisational feature among divergent arenavirus IGRs that is not captured by sequence identity or global RNA-structure similarity. The data are consistent with a model in which a topologically central helical organisation may be associated with conserved regulatory capacity, whereas structurally variable peripheral elements may contribute to lineage- or segment-specific host-factor interaction profiles.

## INTRODUCTION

Biological function is often inferred from sequence conservation, which provides the foundation for most comparative analyses of genes and regulatory elements. For structured RNAs, this relationship is frequently extended to a sequence-structure-function paradigm in which sequence determines RNA fold and fold determines biological activity. However, many functional RNAs challenge this framework. Regulatory RNAs can maintain equivalent functions despite extensive sequence divergence, suggesting that selective pressures may act on organizational features beyond primary sequence, secondary structures or tertiary architectures. Understanding what level of RNA organization remains conserved when sequence similarity or structural contraints are lost remains a central challenge in RNA biology and evolution.

Arenaviruses provide a tractable system in which to address this question. They posses a bi-segmented ambisence RNA genome where each segment, small (S) and large (L), encodes two oppositely oriented open reading frames separated by noncoding intergenic region (IGR). IGRs contain cis-acting signals for transcription termination (Lee et al. 2000), where the viral RNA polymerase recognises specific structural elements to release nascent transcripts. Experimental and computational analyses have shown that IGRs are highly structured, often more than the flanking coding regions, and disruption of their structures can impair viral transcription (López and Franze-Fernández 2007). Yet, comparative analysis reveals that IGRs also exhibit considerable variation in their global architecture (Iwasaki et al. 2016; Kiening et al. 2017) with differences in the number, length, and arrangement of stem-loops between L- and S-segment IGRs, and among Old World (OW) and New World (NW) arenavirus lineages (Kiening et al. 2017). Remarkably, experimental studies have also demonstrated that recombinant arenaviruses with heterologous or synthetic IGRs from distantly related phylogenetic classes can still retain viability and genetic stability (Iwasaki et al. 2016), although often with attenuated phenotypes. These observations suggest that functional conservation is maintained despite extensive sequence and structural divergence and raise the possibility that selective pressures act on organizational features beyond these parameters.

Recent studies of picornavirus internal ribosome entry sites (IRES) have revealed that RNA domains with little detectable sequence homology can nonetheless adopt strikingly similar architecture (Koirala et al. 2019) and employ related mechanisms for host-factor recruitment (Banna et al. 2026). These findings introduce the concept that topology, defined broadly as the connectivity and relative orientations of RNA structural elements (Lescoute and Westhof 2006), may represent an evolutionarily constrained level of RNA organization that persists despite sequence divergence. This principle is also established for viral exoribonuclease-resistant RNAs, in which divergent flaviviral and dianthoviral RNAs converge on pseudoknot-stabilised architectures that encircle the RNA 5′ end and mechanically block 5′-3′ exonucleases (MacFadden et al. 2018; Steckelberg et al. 2018). However, these described systems largely retain recognizable conformations or conserved junction and threading architectures. Whether topology remains informative when global RNA structures become divergent, remains unresolved.

Arenavirus IGRs are uniquely suited to address this question because they combine extensive sequence diversification, considerable structural heterogeneity, and experimental evidence of functional interchangeability. To investigate whether topology explains this apparent paradox, we performed comparative analysis of 58 IGRs spanning the known arenavirus phylogeny using graph-theory based representations (Gan et al. 2004; Kim et al. 2014). Such representations have been previously used to classify RNA structural space (Mallet et al. 2022), identify recurrent topological motifs (Schlick 2018; Kim et al. 2013a, 2013b), and guide RNA design, but rarely been applied to test biological hypotheses concerning functional conservation across natural RNA families. Our analysis revealed that neither sequence nor structural similarity based clustering could recapitulate the functional relationships of IGRs. Instead, a topologically central helical core was persistent across divergent IGR folds wherase topologically peripheral motifs varied across the phylogenetic groups. Functional analysis of a highly peripheral Golden Gate virus (GGV) S-segment IGR further demonstrated selective engagement of antiviral host factors in cells infected with Lymphocytic Choriomeningitis virus (LCMV), a mammarenavirus and whose IGRs did not contain any peripheral elements.

Taken together, by integrating structural and functional perspectives, our findings reveal a likely architectural principle whereby structured viral RNAs achieve functional robustness through conserved topological cores while maintaining evolutionary flexibility through peripheral elements that engage host factors and tolerate variation.

## RESULTS

### Functional conservation of arenavirus IGRs cannot be explained by sequence or structural similarity

The ability of phylogenetically distant arenavirus IGRs to retain equivalent regulatory functions raises the question of whether their evolutionary relationships are reflected at the sequence or structure levels. To address this, we compared all available IGR sequences from mammarenaviruses and reptarenaviruses (Supplementary Information 1) using pairwise similarity measurements and hierarchical clustering. The resulting landscapes revealed inconsistent patterns (**Figure *1***).

**Figure 1:**
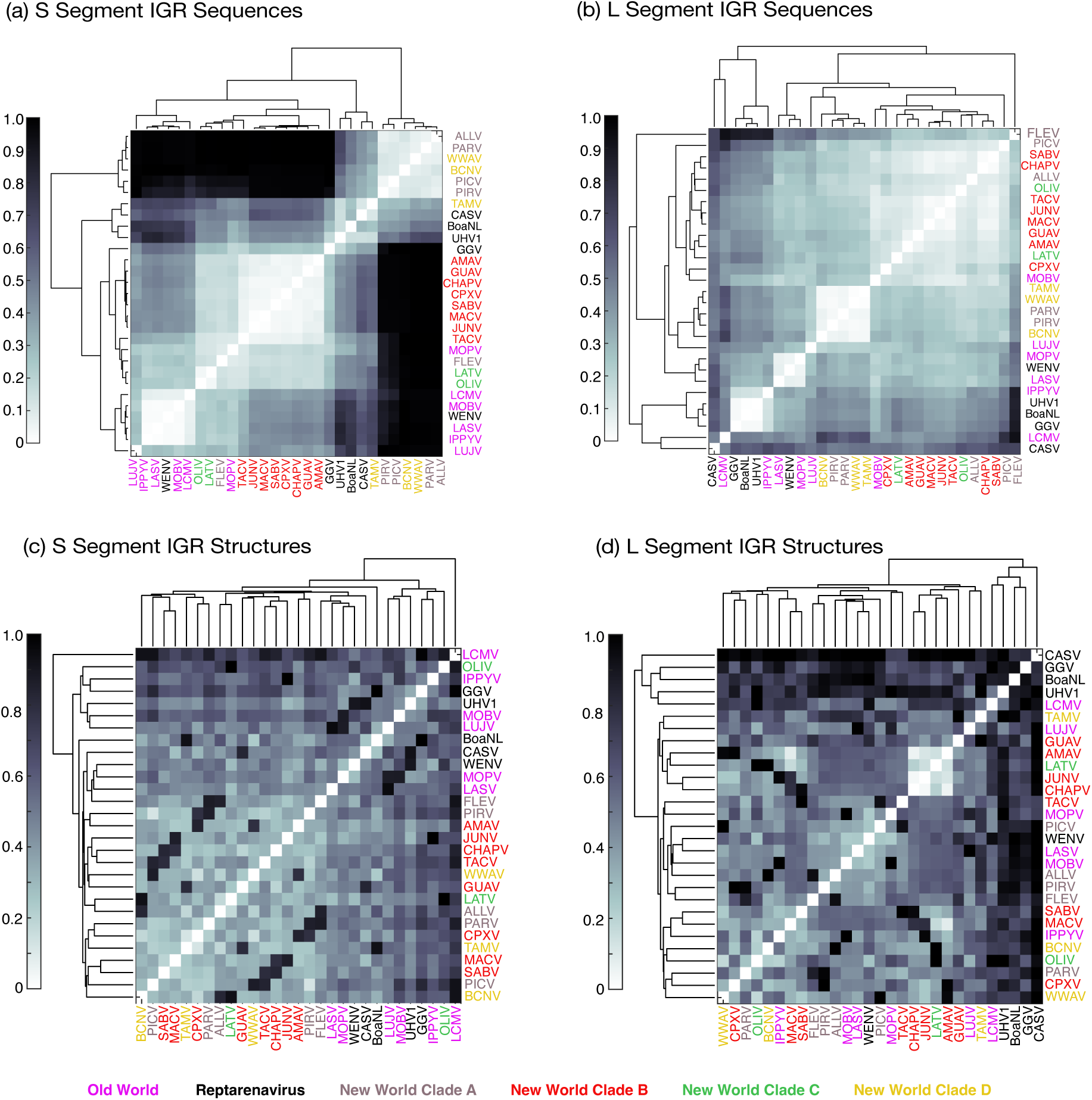
Pairwise similarity analysis of all 58 available arenavirus IGR sequences (a,b) and 2D structures (c,d). Sequences are analysed by clustering of the alignment scores normalised to maximum pairwise distance in the dataset. 2D structure comparison is based on normalised bpRNA-align scores (Lasher and Hendrix 2023). The dendrograms are used here as a visual ordering tool produced by hierarchical clustering rather than as a phylogenetic estimate of evolutionary relationships.

Sequence-based analysis revealed that S-segment IGRs largely segregated according to established arenavirus phylogeny and host range (**Figure *1***a). Notably, New World (NW) mammarenavirus Clade A and D S-segment IGRs formed a distinct cluster that was clearly separated from all other mammarenavirus and reptarenavirus IGRs. In contrast, L-segment IGRs (**Figure *1***b) revealed several relationships that were inconsistent with established arenavirus phylogeny. Most notably, the Old World (OW) L-segment IGRs of Lymphocytic choriomeningitis virus (LCMV) and Ippy virus (IPPYV) clustered with several reptarenavirus IGRs. However, because the clustering metric is derived from pairwise alignment scores and is therefore sensitive to indels events (see Methods), this apparent cross-lineage relationship is likely driven by the unusually large size of LCMV, IPPYV and reptarenavirus IGRs, which are frequently twice the typical length of most arenavirus L-segment IGRs, rather than by genuine sequence conservation. Together, these analyses demonstrate that sequence conservation is unevenly distributed across arenavirus IGRs and may not sufficiently provide a unified explanation for the experimentally observed functional interchangeability of distantly related IGRs.

Arenavirus IGR sequences are constrained by the requirement to form conserved secondary structures that can mediate transcription termination. Despite this structural requirement, structure-based clustering did not reveal simple lineage- or segment-specific groupings. The L-segment IGRs spanned a wide range of secondary structures that did not generally follow any phylogenetic relationships (**Figure *1***d) while the S-segment IGR structures were broadly split across two clusters encompassing mammarenaviruses and reptarenaviruses (**Figure *1***c) and lost any similarity relationships evident at the sequence-level clustering analysis (**Figure *1***a). This structural heterogeneity suggests that functional constraints do not enforce a single conserved, global fold on the IGRs. Instead, arenavirus IGRs may operate within a permissive structural landscape in which multiple distinct RNA architectures are compatible with transcription termination.

### Functionally interchangeable arenavirus IGRs adopt distinct global RNA architectures

To investigate the structural basis of IGR functional conservation, we selected the LCMV and GGV IGRs as representative systems and determined their structures using cryogenic OrbiSIMS (cryo-OrbiSIMS) data restrained modelling (Passarelli et al. 2017; Ward et al. 2024). These IGRs occupy distant positions within arenavirus phylogeny, originate from mammalian and reptilian arenavirus lineages, respectively, and exhibit limited sequence and structural similarities (**Figure *1*** and **Supplementary Information 2**). Despite this evolutionary separation, previous recombinant virus studies demonstrated that the GGV S-IGR can substitute for the LCMV L-IGR while preserving viral viability and genetic stability (Iwasaki et al. 2016, 2015), providing an opportunity to examine how function can be retained across highly divergent RNAs.

Cryo-OrbiSIMS is a highly sensitive solid state mass spectrometry technique for high-resolution chemical and structural analysis of biological material. Previously, we benchmarked this technique (Ward et al. 2024) to demonstrate that the cryo-OrbiSIMS spectrum is a function of biomolecular structure of RNAs and not its chemical sequence, and pseudoenergy restraints derived from the spectral assignments (Borkar 2024) enable experimentally guided modelling of native RNA systems with high precision.

Cryo-OrbiSIMS-guided RNA modelling (Ward et al. 2024) (**Figure *2***) of the L- and S-segment IGRs of the LCMV and GGV revealed that all the four IGRs adopt globally distinct 2D folds with low structural identities (**Supplementary Information 3A**) and marked differences in their organisational complexity (**Figure *2*a-d**). For example, while both LCMV L-IGR and GGV S-IGR contain multiple stem-loop elements, their organisation and global architectures differed substantially, sharing only 2.3% structural identity (as calculated by BEAGLE alignment (Mattei et al. 2015)), which is not significantly greater than that expected for random sequences of comparable lengths (z-score = −0.244 (Mattei et al. 2015)). In the LCMV L-IGR (**Figure *2*a**), the stem-loops radiate from a central four-way junction, forming a radially arranged architecture that includes a basal stem helix (SH) and three stem-loops (SL1–SL3) (Kiening et al. 2017). The LCMV S-IGR (**Figure *2*b**) is considerably simpler, consisting primarily of a single extended helical element, while the GGV L-IGR (**Figure *2*c**), despite being comparable in length to the GGV S-IGR and LCMV L-IGR, is likewise dominated by a single helical segment. Together, these observations demonstrate that substantial architectural diversification occurs not only between arenavirus lineages but also between the L- and S-segment IGRs within individual viruses.

**Figure 2:**
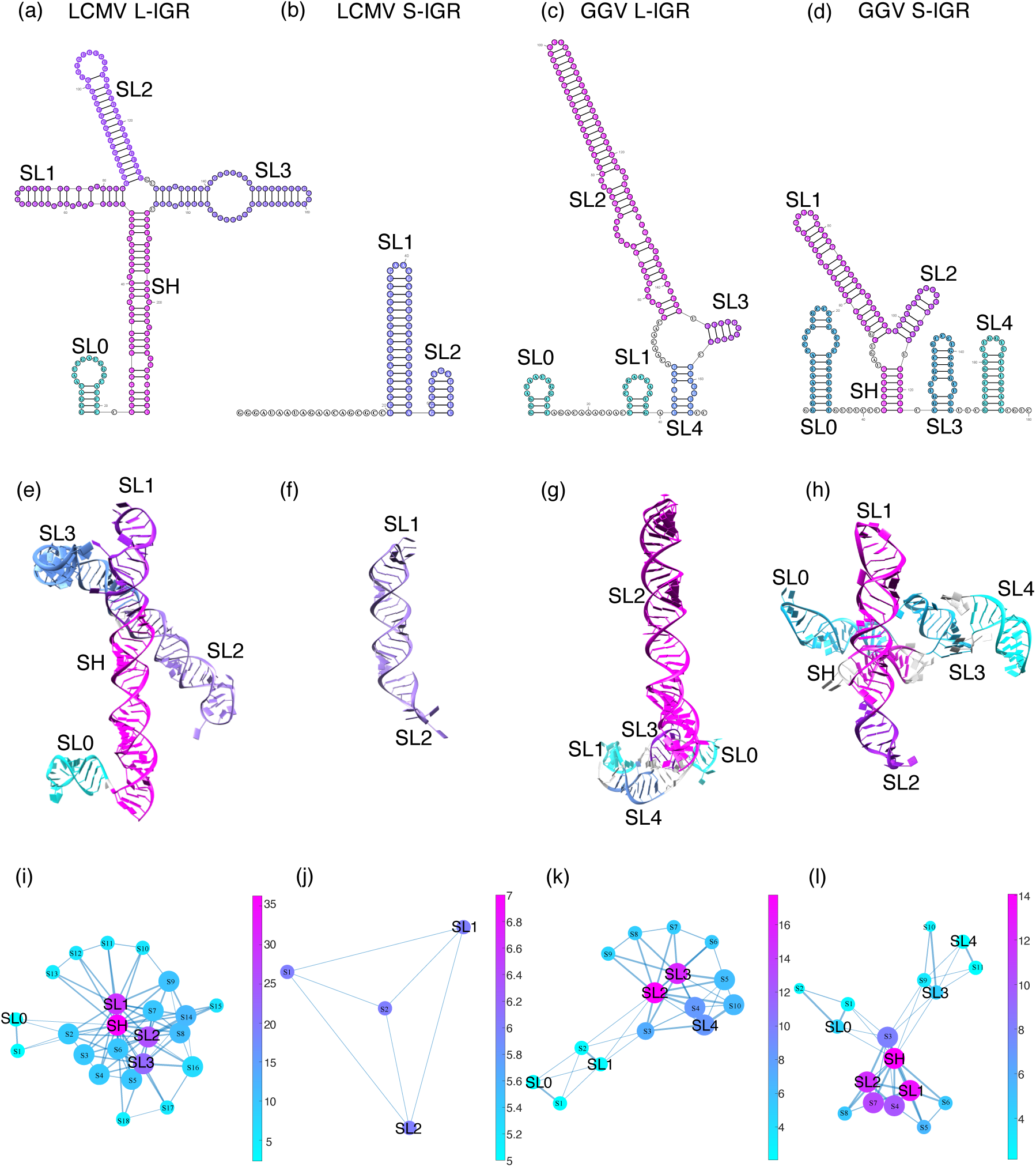
Cryo-OrbiSIMS-guided IGR structures represented as (a-d) secondary structures, (e-h) 3D models and (i-l) topological graphs. In the graphs, each helix is represented as a node, and features such as loops and bulges that connect them are represented as heuristically weighted edges on a scale of 0-1.5 (see Methods for details). The node size is proportional to k-core centrality metric and the node colour is proportional to the strength centrality metric. In 2D and 3D representations, the structural elements are also colour coded according to their node strength for correspondence.

To independently validate the cryo-OrbiSIMS derived models, we performed single-molecule optical tweezer measurements on the LCMV L-IGR and GGV S-IGR, selected as representative IGRs because their multi-stem-loop architectures contain the greatest structural complexity among the four experimentally characterised RNAs. For the LCMV L-IGR (**Figure *3*a**), force-extension measurements revealed three clusters of intermediate states, consistent with four major unfolding events. The contour length changes associated with these transitions are in good agreement with the structural elements determined by the cryo-OrbiSIMS model (**Supplementary Information 4**). Importantly, the calculated contour length of SL1 from the cryo-OrbiSIMS model (18.88 nm) closely matched the experimental value, whereas the corresponding unrestrained prediction from Fold was substantially longer (24.19 nm) (**Supplementary Information 4**).

**Figure 3:**
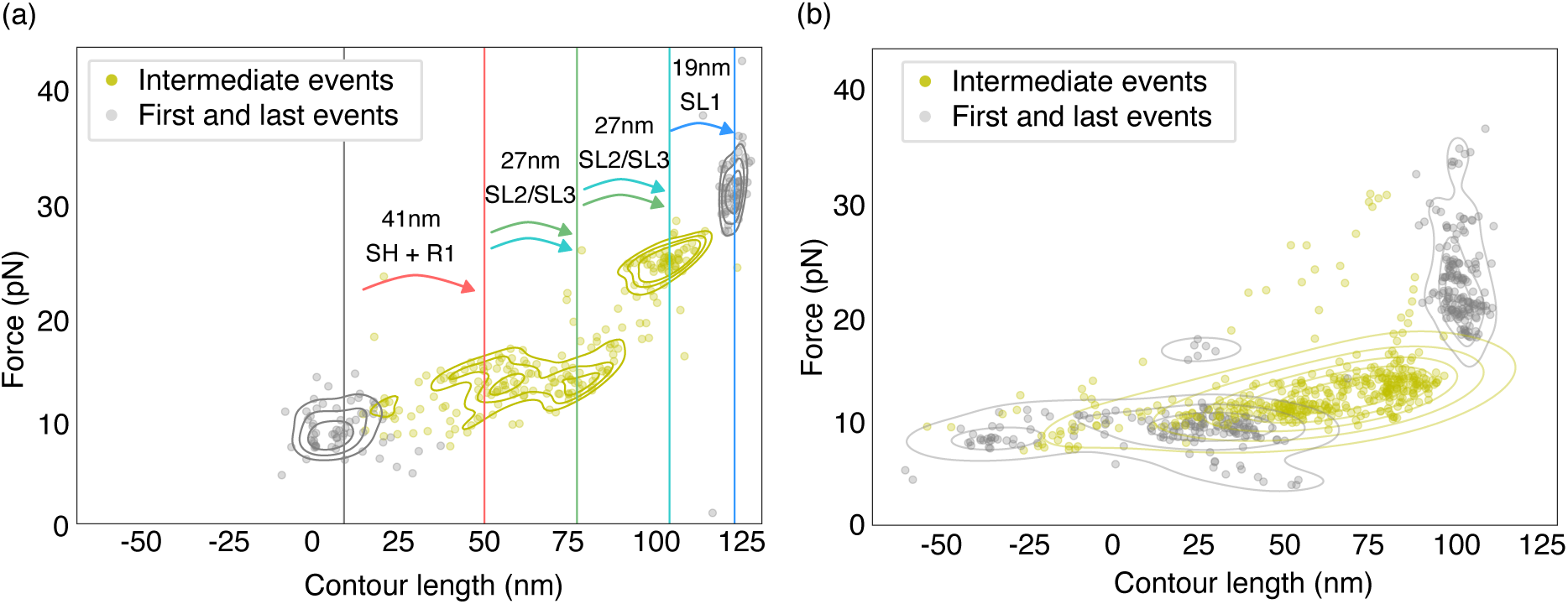
Scatter plot of the observed contour lengths for the individual unfolding events in (a) LCMV L-IGR and (b) GGV S-IGR. The first and last events are coloured grey and olive green indicates intermediate events. Traces were aligned at their fully unfolded length. For LCMV L-IGR, traces were aligned at 122 nm and vertical lines represent transitions that follow unfolding order, which is in turn derived from constant velocity pulling experiments. For GGV S-IGR, traces were aligned at 102 nm. Negative contour lengths may be caused by tertiary interactions.

In contrast to the LCMV L-IGR, the validation of the GGV S-IGR was complicated by the apparent structural heterogeneity of its 5′ region and possible non-native, long-range interactions of the non-native linker sequence introduced within the GGV S-IGR for in optimial in vitro trnascription (see Methods). Cryo-OrbiSIMS and optical tweezers validation was thus performed on a construct without the linker sequence (**Figure *3*b** and **Supplementary Information 5**). Cryo-OrbiSIMS models shared only about 20% structural identity with the *in silico* predictions. Here, the the *in silico* predictions featured a stable stem-loop structure spanning much of the 5’ half of the RNA whereas the cryo-OrbiSIMS models exhibited unpaired or weakly paired structures in this region (**Supplementary Information 5**). Optical tweezers force-extension curves (**Figure *3*b**) yielded a median contour length of approximately 72 nm, substantially shorter than the ~106 nm expected for complete unfolding of the transcript, suggesting that a large proportion of the 5′ sequence exists in an extended or weakly structured state in solution (**Supplementary Information 5a)**. Thus, although unlike LCMV L-IGR, direct structural mapping between cryo-OrbiSIMS and optimal tweezers data was not possible due to the structural heterogeneity of GGV S-IGR in solution, structural observables from both these techniques are more consistent with each other than with the *in silico* predictions. Taken together, these results indicate that the cryo-OrbiSIMS-derived architectures are representative of the predominant conformations adopted by both LCMV L- and GGV S-IGRs in solution.

Using these validated cryo-OrbiSIMS-derived secondary structures, we next modelled the three-dimensional conformations of the IGRs using RNAComposer (Ward et al. 2024; Popenda et al. 2012) (**Figure *2*e-h**). Consistent with the 2D analyses, the resulting 3D models revealed pronounced global divergence between the IGRs (**Supplementary Information 6**). The LCMV L-IGR adopts an extended X-shaped topology (**Figure *2*e**), characterised by coaxial stacking of SL1 over the SH (Lescoute and Westhof 2006), while SL2 and SL3 also stack coaxially over one another and at an acute angle relative to the SL1–SH axis forming a compact, X-shaped structure. In contrast, the GGV S-IGR maintains a predominantly linear topology (**Figure *2*h**), in which the two branches of SL1 and SL2 stack coaxially, whereas SL3 and SL4 remain structurally independent. The LCMV S-IGR and GGV L-IGR similarly retain their dominant stem-loop element as an extended helical segment in the corresponding 3D structures (**Figure *2*f,g**).

### A graph-defined central helical organisation is recurrent among divergent arenavirus IGR folds

Despite the extensive diversification observed across arenavirus IGRs (**Figure *1***) and the pronounced architectural differences revealed by experimental structure determination (**Figure *2***), the LCMV and GGV IGR structures seemed to contain a common organisational feature: in all four experimentally characterised IGRs, one or more helical elements occupied structurally dominant position within the fold, frequently participating in coaxial stacking interactions despite substantial differences in global architecture. We therefore asked whether topology, rather than sequence or 2D-fold similarity, provides a recurrent organisational framework across arenavirus IGRs.

To address this question, we represented the IGR topology as graphs (Gan et al. 2004; Kim et al. 2014) (**Supplementary Information 8**). Here, helical segments were represented as graph nodes, and features such as loops, bulges and junctions were defined as the edges connecting them. The graph edges were also heuristically weighted on whether they would aid coaxial stacking of the nodes, allowing the global organisation of each RNA to be quantitatively assessed for presence of structurally central coaxial stacks.

Application of this framework to the experimentally determined IGR structures revealed that the dominant helical elements identified in the three-dimensional models consistently occupied topologically central positions within their corresponding graphs (***Figure 2***i-l). In the LCMV L-IGR, the basal helix (SH) and adjacent stem-loops SL1-SL3 formed a densely interconnected topological core, consistent with the compact four-way-junction architecture observed in the structural model. Similarly, the coaxially stacked SL1-SL2 module of the GGV S-IGR occupied the most central position within the graph despite its markedly different overall architecture with the LCMV L-IGR. In contrast, SL0 was identified as peripheral in each IGR. SL0 originates from linker residues introduced during our *in vitro* transcription template design and is structurally isolated from the native IGR architecture, it thus provides an internal validation of the graph-based approach.

Extending the graph-based analysis to all available IGRs from 29 distinct mammarenavirus and reptarenaviruses (**Supplementary Information 1**), we found that nearly all L- and S-IGRs, irrespective of length, sequence identity, segment origin or phylogenetic lineage, had high graph densities (**Supplementary Information 8a-b**) and at least one highly connected central node (**Supplementary Information 8c**) that represented a dominal helical element in the respective IGR. A total of 67 of 148 (45%) of such helical segments were classified as central, whereas 27 (18%) were identified as topologically peripheral in 23 distinct IGRs (**Figure *4***a). Peripheral segments were particularly abundant (≥70% of all identified segments) in both L- and S-IGRs of reptarenaviruses, as well as in L-IGRs of OW arenaviruses and NW Clade C viruses (**Figure *4***b). In contrast, NW Clade A and B viruses exhibited a marked asymmetry between genome segments, with S-IGRs containing substantially more peripheral elements than their corresponding L-IGRs (**Figure *4***b). Thus, within the analysed dataset, IGR graphs consistently contained one or more highly central helical elements, whereas the abundance and placement of peripheral elements varied between segments and phylogenetic groups. These observations identify a recurrent organisational pattern that is compatible with candidate conserved core functions and greater structural flexibility at peripheral nodes.

**Figure 4:**
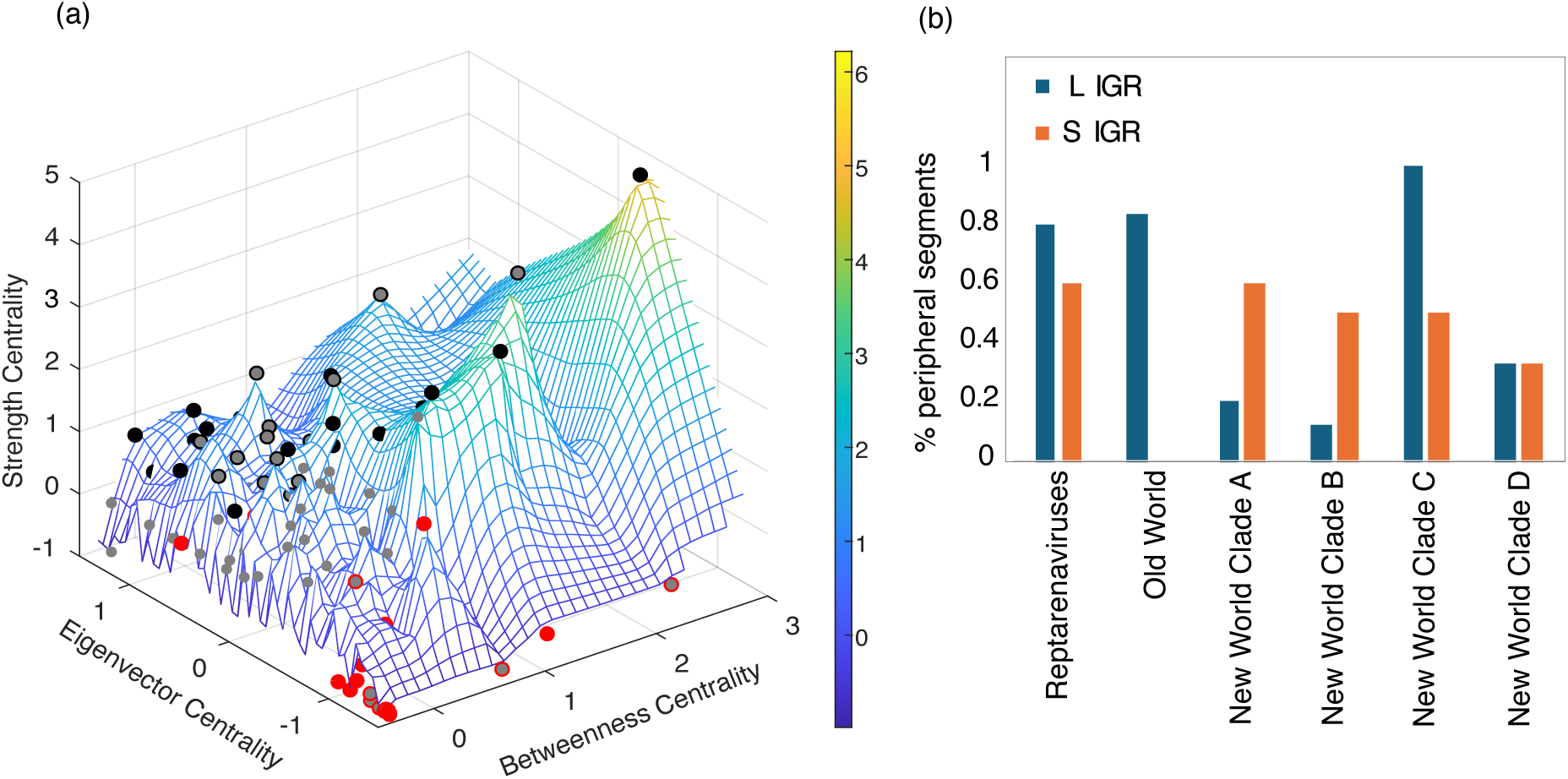
Graph-based analysis of IGRs. (a) Distribution of graph centrality metrics: strength, eigenvector and betweenness, for the 58 IGR RNA graphs consisting of 148 individual stem-loop segments from 29 different arenaviruses (Supplementary Information 1). All values are standardised to z-scores (mean = 0 and standard deviation = 1) for plotting and individual axes are trimmed to the limits shown in this figure for easier data visualisation. Positions of the IGR nodes (see Supplementary Information 7) are marked on the surface as dots. Central nodes are denoted as solid black and peripheral nodes are denoted as solid red dots. Any unclassified nodes are termed “neutral” and indicated as grey. (b) Abundance of topologically peripheral segments within phylogenetic classes of arenaviruses.

To ascertain if this classification of central and peripheral was dependent on the heuristic edge weights used, we also randomly perturned the weights by 20% over 100 iterations. The centrality and peripherality measures were observed to be stable (**Supplementary Information 9a**) under this level of uncertainty. Across the perturbations, 64 of 67 originally central nodes (95.5%) and 17 of 27 peripheral nodes (63.0%) retained their classification in ≥90% of iterations (**Supplementary Information 9b-c)**, Secondly, we aligned the graphs by establishing node correspondence using their degree, eigenvector centrality, and betweenness centrality metrics. Amongst all the aligned nodes, more than 93% nodes which were classfied as central segment in one IGR, also aligned to a central segment in another IGR (**Supplementary Information 10)**. This high alignment fidelity of central nodes underscores their structural indispensability, while the moderate stability of peripheral nodes may reflect their context-dependent roles. Lastly, as an external comparator, we applied the same graph framework to flavivirus 3′ UTR RNA structures previously derived from an integrated chemical probing and computational modelling approach (Villordo et al. 2015). Here conserved pseudoknot-based architectures are not expected to be dominated by coaxially stacked helices. In contrast to arenavirus IGRs, node centrality values were substantially less stable following perturbation of edge weights (**Supplementary Information 9d**), indicating that the robust central helical organisation observed for arenavirus IGRs is not an automatic consequence of applying the framework to structured viral RNAs.

### Topologically peripheral IGR elements are associated with distinct host-factor interaction profiles

The observation that most topological variation among arenavirus IGRs is concentrated within peripheral stem-loop elements prompted us to ask whether these structures contribute to any biological functions beyond maintenance of core IGR activity. We therefore focused on the GGV S-segment IGR, which contained the highest abundance of peripheral elements within the entire arenavirus IGR dataset (**Supplementary Information 8c**), and compared its host interaction landscape with that of the LCMV L-IGR, whose topology is almost entirely dominated by the central helical core (**Supplementary Information 8c**).

To identify cellular factors associated with these structurally distinct IGR architectures, biotinylated GGV S-IGR and LCMV L-IGR RNAs were used as affinity baits during rLCMV-eGFP infection, followed by quantitative proteomic analysis of recovered host proteins (**Figure 5**a). A total of 65 differentially enriched proteins were identified for GGV S-IGR compared to 35 for LCMV L-IGR (**Supplementary Information 11**). Gene Ontology analysis revealed marked differences between the two interactomes (**Figure 5**b). Proteins associated with the LCMV L-IGR were predominantly enriched for general RNA metabolism, translation, and gene-expression pathways. In contrast, the GGV S-IGR interactome additionally exhibited strong enrichment for pathways associated with antiviral responses (GO:0050792), viral genome regulation (GO:0045069), and host defence mechanisms (GO:0051607). This enrichment was driven by several canonical antiviral RNA-binding proteins, including IFIT1, PKR (EIF2AK2), ISG15, ADAR1 (DSRAD), ILF3, DDX21 and DHX15 (**Figure 5**c). Notably, analysis of raw abundance profiles for GGV S-IGR interacting proteins across infected and mock-infected backgrounds (**Figure 5**c) revealed that IFIT1, PKR and ISG15 were selectively detected in GGV S-IGR pull-downs obtained from infected cells and were absent from both the negative control and the LCMV L-IGR interactome. This indicates that these proteins have a GGV S-IGR–specific association that is observed only in the context of infection. However, since global abundance of these proteins was not independently assessed, this scenario potentially reflects either specific induction of these proteins upon infection or their recruitment to the IGR via infection-induced cofactors. Other proteins, including ADAR1, ILF3 and DHX15, were recovered from both infected and mock-infected backgrounds but remained enriched relative to control pull-downs, suggesting stable interaction with the GGV S-IGR independent of infection status. Collectively, these findings indicate that the GGV S-IGR engages a distinct host-factor repertoire enriched for proteins involved in antiviral RNA sensing, RNA editing and translational regulation.

**Figure 5:**
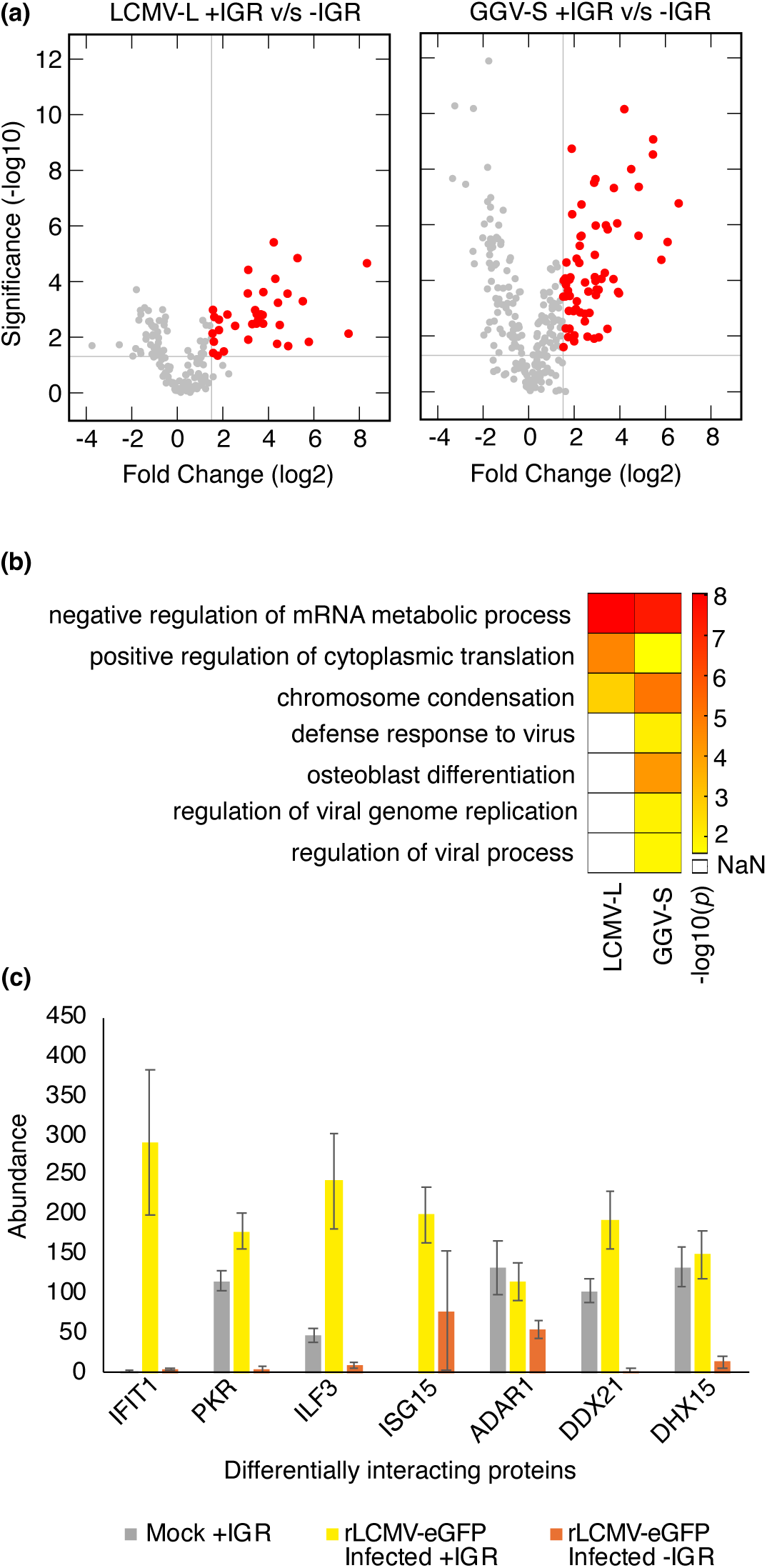
Proteomics analysis of IGR-host interactome. (a) Volcano plots for LCMV L-IGR and GGV S-IGR pull-down from rLCMV-eGFP infected cells and normalised against negative control (no IGR present in the pull-down experiment). (b) Semantically reduced GO enrichment categories for the significant hits show that LCMV L-IGR interactome is enriched for generic RNA metabolism and gene expression factors, whereas GGV S-IGR interactome additionally contains pathways linked to virus regulation. (c) GGV S-IGR specific GO categories are driven by specific innate antiviral factors. Raw abundance values for these proteins are indicated for GGV S-IGR pull-down experiments across rLCMV-eGFP infected and mock backgrounds as well as a no-IGR control. Mean values are plots across six replicates (including 3 biological x 2 technical replicates) for each condition, error bars denote standard deviation.

To determine whether these host factor interactions are specifically mediated by topologically peripheral elements in the GGV S-IGR (**Figure *2***h,l), we compared cryo-OrbiSIMS fragmentation profiles of GGV S-IGR in its apo state and following host-protein capture. Although the global assignment profiles remained highly correlated (**Figure *6***a, Pearson R = 0.97), indicating preservation of the overall RNA architecture, localized changes were concentrated within the peripheral SL0, SL3 and SL4 stem-loops (**Figure *6***b). These regions displayed a significant enrichment of residues with large Δz values (hypergeometric p = 0.00046) (**Figure *6*b**), where Δz quantifies the change in standardized cryo-OrbiSIMS assignment profiles between protein-bound and apo RNA states.

**Figure 6:**
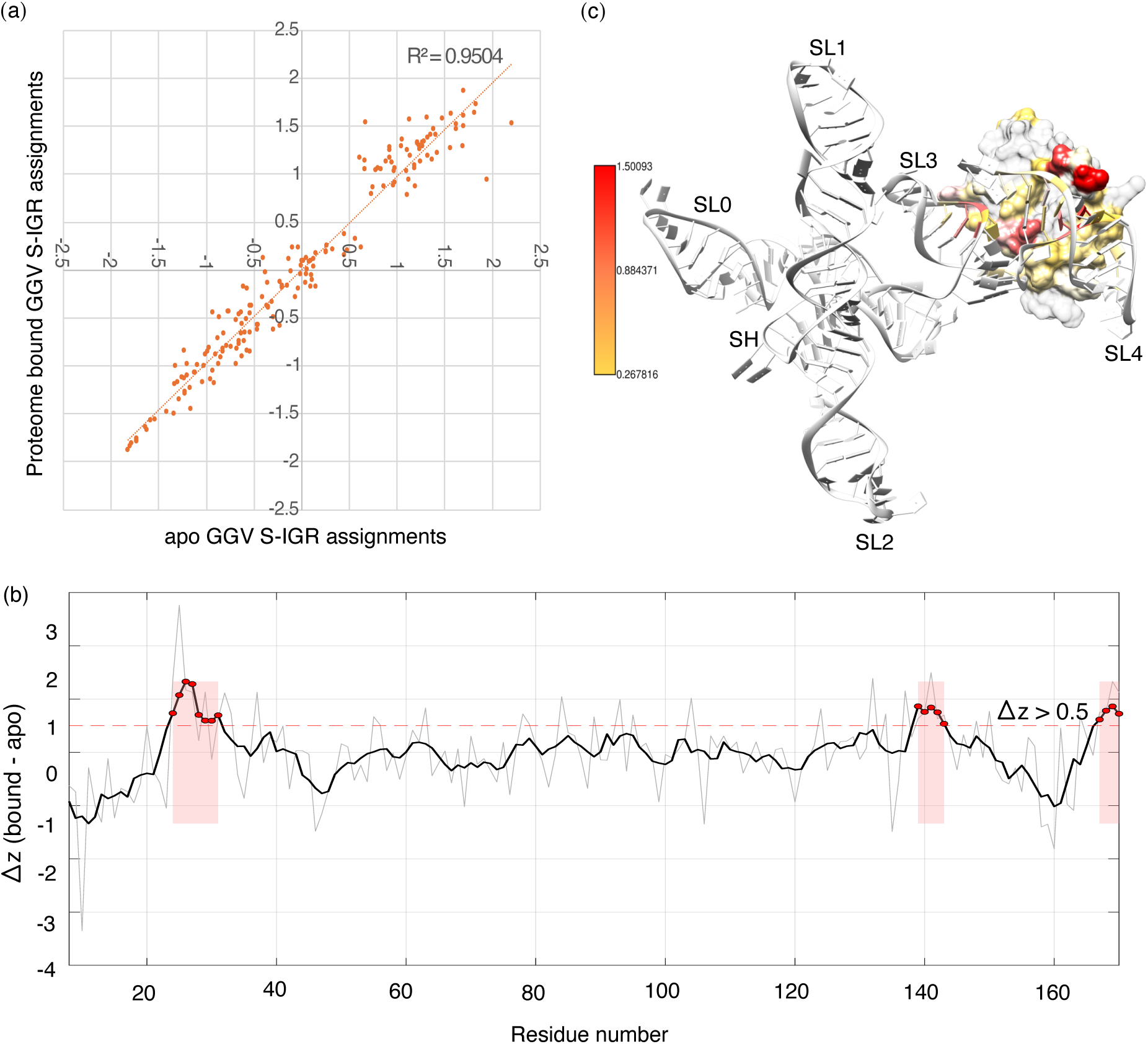
(a) Comparison of cryo-OrbiSIMS assignments for GGV S-IGR in the pull-down protein bound samples v/s apo RNA samples. The assignments were standardised to z-score and averaged over a sliding window of 4 nucleotides to remove noisy fluctuations from per residue assignments. (b) Distribution of difference (Δz) between bound and apo z-scores across the full-length IGR. Dotted red line indicates the Δz cut-off of 1, all data points with Δz>1 are indicated as red dots and the corresponding region along the IGR sequence is shaded red rectangle. (c) Docking of the double-stranded RNA binding motif of PKR (dsRBM PDB ID 1QU6), one of the top-two hits identified from our proteomics analysis, on the cryo-OrbiSIMS model of GGV S-IGR. For the docked structure, the colour bar indicates log2(fold change) of the number of native contacts of the RNA and protein in the docked state compared to their respective apo forms.

The localisation of the observed cryo-OrbiSIMS assignment differences to peripheral stem-loops is consistent with local structural perturbations in regions associated with the GGV S-IGR pull-down profile. However, these data do not establish whether the altered assignments arise from direct protein binding, indirect effects of protein complexes, or changes in the RNA conformational ensemble. Among the enriched proteins, PKR was particularly notable because it contains a well-characterized double-stranded RNA-binding domain (Spanggord et al. 2002), raising the possibility that the peripheral elements of the GGV S-IGR provides a structurally accessible motif for antiviral protein recognition. We therefore performed docking simulations in HADDOCK (Dominguez et al. 2003) using the NMR structure of PKR RNA-binding domain (PDB ID 1QU6) and the cryo-OrbiSIMS GGV S-IGR model (**Figure *6***c). These consistently positioned PKR adjacent to the peripheral SL3-SL4 region in agreement with both the proteomic enrichment of PKR and the localised Δz perturbations observed in this region of the IGR in the pull-down samples. Although these docking models do not establish *in vivo* binding or binding specificity, they support the interpretation that SL3-SL4 may provide structurally accessible motifs capable of engaging dsRNA-binding host factors. Taken together, these results nominate topologically peripheral elements as candidate regions through which divergent IGRs may engage distinct cellular interaction environments. Targeted deletion and compensatory-structure experiments will be required to establish the contribution of individual peripheral elements to protein recruitment and IGR function.

## DISCUSSION

A longstanding assumption in RNA biology is that functional conservation arises through preservation of sequence and, where sequence diverges, through preservation of structure. Arenavirus IGRs challenge this framework because these experimentally interchangeable regulatory RNAs exhibit extensive divergence at both levels. If the IGRs shared closely related folds, their functional compatibility would be unsurprising and could be explained through conventional models of structural conservation. Instead, the LCMV and GGV IGRs occupy distant regions of structural space yet retain equivalent regulatory capacity in recombinant viruses. Functional conservation in this context therefore becomes non-trivial and raises the question of what organizational feature is preserved. By integrating phylogenetic comparison, experimentally informed RNA modelling and graph-based analysis, we identify RNA topology as a candidate conserved organisational layer among the arenavirus IGRs examined. While sequence similarity and global RNA architecture vary extensively across arenavirus IGRs, each RNA retains a highly connected and central helical organization containing one or more stem-loop elements that persists across divergent folds. Such behaviour is consistent with RNA evolutionary theory, wherein functional RNAs are thought to occupy expansive neutral networks in sequence and structure space (Schuster et al. 1994), allowing extensive divergence in global fold while preserving core functional outputs (Schultes and Bartel 2000). The ability of divergent IGR architectures to support viable recombinant viruses therefore reflects the inherent robustness and modularity of regulatory RNA elements (Kim et al. 2014).

Our findings extend previous observations that higher-order RNA architecture can be conserved despite sequence divergence. Structural studies of picornaviral IRES elements revealed common architectural motifs and topological arrangements across distinct viral RNA classes despite limited sequence homology (Koirala et al. 2019; Banna et al. 2026). In comoviruses, stem-loop elements SL1 and SLC residing on separate genomic segments share minimal sequence identity yet are functionally interchangeable in supporting RNA accumulation, and separate exchanges of their stem and loop portions are tolerated without loss of function (Guo et al. 2017). Similarly, internal ribosome entry sites (IRES) from different picornaviruses adopt distinct secondary and tertiary folds but all recruit host translational machinery to initiate cap-independent translation, demonstrating convergent functional outcomes across structurally diverse motifs (Fernández-Miragall et al. 2009). In flaviviruses, exoribonuclease-resistant RNAs (xrRNAs) across genera fold into different pseudoknot configurations, yet universally block host exonucleases to produce subgenomic RNAs critical for replication and pathogenicity, again highlighting conserved function amid structural variation (Villordo et al. 2015). Collectively, these examples demonstrate that equivalent biological functions can be preserved despite substantial sequence and structural divergence. However, the organizational principles that enable such conservation remain poorly defined. In some systems, conservation is associated with preservation of recognizable architectural motifs, whereas in others, similar functions emerge from markedly different RNA folds. Our findings suggest that arenavirus IGRs reconcile these observations through conservation of a common topological organization with phenotypic consequences.

Several peripheral elements observed in our study seem to occur in reassorting or recombinant arenaviruses. This includes reptarenaviruses, known to undergo frequent reassortments (Lintala et al. 2025; Stenglein et al. 2015), laboratory-generated reassortant ML29 (Carrion et al. 2007) of Lassa virus S- and Mopeia virus L-segment, and Xapuri virus, proposed to be a natural reassortant with S segment closely related to Oliveros virus (Fernandes et al. 2018). The presence of a peripheral element within the Tamiami virus (TAMV) S-IGR is notable in light of reports that this segment arose through recombination between NW clade B and clade A arenaviruses while retaining a NW clade A-like IGR. Similarly, historical recombination events (Archer and Rico-Hesse 2002) are thought to have given rise to the ancestral S segment of the clade containing Sabiá virus (NW Clade B), which we observe to contain significantly higher enrichment of peripheral segments compared to in the L segment and compared to most other arenavirus phylogenetic groups. While these associations are consistent with a role in tolerance for sequence and structural variability or adaptive flexibility, we acknowledge that in the remit of the current study, they remain correlative and do not establish causality. Individual elements will thus require future targeted experimental interrogation to determine their selective functional significance.

In summary, our findings indicate that functional conservation among arenavirus IGRs is not adequately explained by sequence identity or preservation of a common RNA fold. Instead, comparison across sequence, structural and graph-based representations identifies a recurrent central helical organisation across the analysed IGRs despite extensive sequence and architectural diversification. Variation was more frequently observed in topologically peripheral elements, which were associated with distinct host-factor interaction profiles in the experimental systems examined. Whether these peripheral elements directly determine host-factor recruitment or functional diversification remains to be tested. Together, these observations suggest that topology represents an important and previously underappreciated level of RNA organization, providing a potential explanation for how regulatory RNAs maintain conserved functions despite substantial evolutionary divergence.

## MATERIALS AND METHODS

### RNA preparation

Intergenic region sequences were obtained from NCBI GenBank (**Supplementary Information 1**) and DNA templates for *in vitro* transcription (IVT) of LCMV and GGV IGRs were chemically synthesised (GenScript UK) with a T7 promoter and short linker sequence (ATAAT) (Conrad et al. 2020) positioned upstream of the sequence of interest cloned into a pUC19 backbone using XbaI and BamHI (or SalI). IVT was performed from linearised pUC19-IGR template using Promega T7 RiboMAX™ Express Large Scale RNA Production System Kit (Catalogue Number P1300) according to the manufacturer’s protocol. The resulting RNA product was purified and concentrated using the Monarch® RNA Cleanup Kit ( (NEB, T2040L) according to the manufacturer’s protocol and transcript quality was assessed using an Aligent: High Sensitivity RNA ScreenTape®, performed at the DeepSeq Facility, Queen’s Medical Centre, University of Nottingham.

### Cell culture, infection and proteomics

Human lung adenocarcinoma epithelial cells (A549 cells) were infected with recombinant rLCMV-eGFP as previously described (Shaw et al. 2024). Cells were harvested at 72 hours post infection and lysed in ice-cold modified Pierce™ IP Lysis Buffer (Thermo Scientific, 87787) containing 25 mM Tris-HCl pH 7.4, 150 mM NaCl, 1 mM EDTA, 1% NP-40 and 5% glycerol and Halt™ Protease Inhibitor Cocktail (78429). Following 45 minutes incubation at 4°C with gentle agitation, lysates were clarified by centrifugation at 13,000xg for 10 minutes at 4°C, and supernatants collected for analysis. Thermo Scientific™ Pierce™ Magnetic RNA-Protein Pull-Down Kit (20164) was used for the pulldown reactions with bait LCMV L-IGR and GGV S-IGR modified with 3’-desthiobiotinylated cytidine bisphosphate using T4 RNA ligase following manufacturer guidelines. Biotinylated RNA was immobilised on Pierce™ Streptavidin Magnetic Beads, incubated with the clarified cell lysate for 60 min at room temperature with gentle rotation before washing three times with 1x wash buffer (Pierce) adjusted to 150mM NaCl. Beads were resuspended in 50 µl of 50 mM ammonium bicarbonate and flash frozen in liquid nitrogen for proteomics analysis. Mock infected cells and pull-downs in the absence iof IGR RNAs were used as negative controls.

Proteomics was performed at the Advanced Mass Spectrometry Facility, School of Biosciences, University of Birmingham. Data-dependent scanning acquisition was conducted using the Xcalibur 4.0 software. Mass spectrometer scans alternated between full FT-MS scans (m/z 360 – 1600) and subsequent high energy collision dissociation (HCD) MS/MS scans of the 20 most abundant ions. Survey scans were acquired in the Q-Exactive HF with a resolution of 120000 at m/z 200 and an automatic gain control of 3×106. Precursor ions were fragmented in HCD MS/MS with resolution set at 15,000 and a normalized collision energy of 28. The ACG target for HCD MS/MS was 1×105. The precursor isolation window width was 1.2 m/z and only multiply charged precursor ions were selected for MS/MS. Spectra were acquired for 56 minutes with dynamic exclusion time of 20s. A total of three biological replicates were analysed, each measured with two technical injections.

Differential abundance between each bait pulldown and its corresponding negative control was log2-transformed and assessed using a statistical t-test. Proteins were considered significantly enriched if they exceeded an absolute log2FC threshold of 1.5 and an FDR-adjusted p-value below 0.05. All significant hits were selected for downstream functional enrichment analysis using g:Profiler (Kolberg et al. 2023), with FDR correction applied to enrichment p-values. To reduce redundancy among the enriched GO terms, results were processed with REVIGO (Supek et al. 2011) using default parameters (SimRel similarity measure, medium similarity threshold of 0.7). Enriched terms were then filtered to a database frequency of 0-2% to exclude overly general categories, and the remaining terms were sorted by dispensability, with the five least dispensable (i.e., most representative) terms selected for reporting and cross-condition comparison (**Supplementary Information 12**).

### Cryogenic OrbiSIMS

Cryogenic OrbiSIMS measurements were performed as described previously (Ward et al. 2024). Briefly, 100 nM IGR RNA in 1x PBS supplemented with 7.5% glycerol and 5 mM MgCl_2_ was analysed under frozen-hydrated conditions at the OrbiSIMS instrument facility at Nanoscale and Microscale Centre, University of Nottingham UK. For each RNA, three independent technical replicates were collected in negative polarity and buffer alone data was used as reference spectrum. Peak lists were extracted from the raw spectra using IonToF SurfaceLab 7 and processed and assigned using custom scripts implemented in MATLAB R2021a and BASH shell as previously decribed (Borkar 2024; Ward et al. 2024), generating residue-wise assignment counts, which were converted into pseudo-energy restraints for secondary structure modelling using Fold algorithm within RNAstructure (Bellaousov et al. 2013). The resulting restrained secondary structures were subsequently used as inputs for three-dimensional RNA modelling with RNAcomposer (Antczak et al. 2017). Residue-level assignment counts were also used to compare IGR structures in the apo RNA samples and protein-enriched pull-down samples. Assignment counts were standardised to z-scores, and differences between apo and pull-down samples (Δz) were calculated for each residue. To reduce local variability and highlight regional effects, Δz values were averaged using a sliding window of four consecutive nucleotides. Regions displaying pronounced deviations from the apo state were interpreted as significant hotspots of local structural changes with positive Δz indicating destabilisation and negative Δz indicating protection by proteins.

### Topological representation

IGR 2D structures were parsed into bpRNA-align format (Lasher and Hendrix 2023) and the structural information converted into graph-based topological networks (**Supplementary Information 8**). Helices identified in the bpRNA-align annotation and the stem-loops containing them were coded as nodes and the structural features connecting them, such as bulges and loops, were coded as edges. The edges were heuristically weighted according to whether they would aid coaxial stacking of the two helices they connected. For example, hairpin loops are weighted 0 as they would cap a helix and not aid in coaxial stack, whereas bulges and internal loops are weighted 1.5 as they are most likely to aid stacking. This weighting scheme allowed the graph to encode not only connectivity but also the likelihood of helix-helix stacking interactions, enabling identification of stacked structural motifs from the RNA topology alone. The edges were also inversely penalised for length, as shorter connectors are likely to lead to structural stability.

Graph centrality metrics such as strength, eigenvector centrality and betweenness centrality to quantify the topological properties of indidivual nodes. Strength was defined as the sum of the weights of all edges incident on a node. Eigenvector centrality was calculated using edge weights as measures of connection importance, whereas betweenness centrality was calculated using inverse edge weights as path costs. Betweenness values were log-transformed, and all measures were standardised to z-scores within each graph. To account for the dependence of centrality measures on node strength, linear models were fitted between strength and eigenvector centrality, and between strength and betweenness centrality; residuals from these models described whether a node had greater or lower strength than expected for its centrality values. Nodes were classified as central when they had high strength (z>1) and positive residuals in either the strength-eigenvector or strength-betweenness relationship. Nodes were classified as peripheral when they had below-average eigenvector centrality and a negative residual in the strength-betweenness relationship. Nodes that did not meet either criterion were classified as neutral.

To assess the robustness of the central/peripheral classification with respect to uncertainty in the heuristic edge weights, random perturbation were performed over 100 iterations. In each iteration, edge weights (ranging from 0 to 1.5) were adjusted by up to 20% of their original values. Stability was quantified by the fraction of nodes retaining their original classification in ≥90% of iterations.

### Optical tweezers

The IGR RNA to be analysed was attached to two micro-sized beads using DNA RNA hybrid handles (4kb or 2.5kb in length) as described previously (Pekarek et al. 2022). Beads were caught in two lasers, one mobile and one static, after which the mobile laser slowly moved away at a constant velocity, applying force on the RNA structure. The applied force was used to measure and identify events where structural elements unfolded or dissociated, measuring both the increase in length and the force required for the rupture of the involved interactions. These experiments were repeated multiple times for a given molecule and multiple pairs of beads were measured. The resulting data was analysed using the worm-like chain (WLC) model and the extensible worm-like chain model (eWLC) to obtain molecule length changes upon dissociation or unfolding.

## ACKNOWLEDGEMENTS

The authors would like to thank the School of Veterinary Medicine and Science, University of Nottingham funding to AC, Anne McLaren Fellowship granted by the University of Nottingham to ANB, and the Engineering and Physical Sciences Research Council (EPSRC) grant code EP/P029868/1, which funded the OrbiSIMS instrument at University of Nottingham and the MRC New Investigator Grant MR/W002167/1 awarded to TLF. The authors would like to thank Professor John Barr, University of Leeds, UK for the plasmid constructs used for generating rLCMV-eGFP virus used in this study. The authors would also like to thank Dr Anna Kotowska, Research Fellow and SIMS Facility Manager, nmRC University of Nottingham for performing OrbiSIMS data collection required for this work.

## DATA AVAILABILITY STATEMENT

Raw and processed cryogenic OrbiSIMS data, including peak lists and the derived psuedoenergy terms used for RNA structure modelling are available upon request from the corresponding author. All scripts used in the processing of the cryo-OrbiSIMS data is available at the GitHub repository https://doi.org/10.5281/zenodo.10960751. IGR models in bpRNA-align or graph formats can be made available upon request from the corresponding author. The mass spectrometry proteomics data have been deposited to the ProteomeXchange Consortium via the PRIDE partner repository with the dataset identifier PXD084083. All other data supporting the current study is available through the main text and supplementary information accompanying this article.

