## Supplementary Information for "Comparative analysis identifies a shared topological organisation among divergent arenavirus intergenic RNAs"

Department of Biochemistry, Cell and Systems Biology, Biosciences Building, Crown Street, Liverpool, L69 7ZB

**Supplementary Information 1:** GenBank ID for the IGR sequences analysed in this work

| Virus name | Acronym | ID | Co-ordinate start | Co-ordinate end |
|---|---|---|---|---|
| Lassa_virus | LASV-L | LT601601.1 | 405 | 504 |
| Lujo_virus | LUJV-L | NC_012777.1 | 334 | 437 |
| Mopeia_virus | MOPV-L | NC_006574.1 | 392 | 501 |
| Mobala_virus | MOBV-L | DQ328876.1 | 417 | 512 |
| Ippy_virus | IPPYV-L | NC_007906.1 | 455 | 618 |
| Junin_virus | JUNV-L | KU978799.1 | 395 | 477 |
| Machupo_virus | MACV-L | NC_005079.1 | 383 | 464 |
| Chapare_virus | CHAPV-L | EU260464.1 | 370 | 441 |
| Sabiá_virus | SABV-L | AY358026.2 | 378 | 447 |
| Tacaribe_virus | TACV-L | MT081317.1 | 358 | 440 |
| Pichinde_virus | PICV-L | NC_006439.1 | 6604 | 6658 |
| Whitewater_Arroyo_virus | WWAV-L | NC_010703.1 | 366 | 442 |
| Amapari_virus | AMAV-L | NC_010251.1 | 6613 | 6702 |
| Cupixi_virus | CPXV-L | AY216519.2 | 6658 | 6747 |
| Pirital_virus | PIRV-L | NC_005897.1 | 352 | 426 |
| Paraná_virus | PARV-L | NC_010761.1 | 370 | 450 |
| Flexal_virus | FLEV-L | NC_010759.1 | 339 | 381 |
| Allpahuayo_virus | ALLV-L | NC_010249.1 | 6637 | 6709 |
| Bear_Canyon_virus | BCNV-L | NC_010255.1 | 349 | 421 |
| Latino_virus | LATV-L | NC_010760.1 | 344 | 427 |
| Oliveros_virus | OLIV-L | NC_010250.1 | 6736 | 6816 |
| Lymphocytic_choriomeningitis_virus | LCMV-L | NC_077806.1 | 362 | 563 |
| Golden_gate_virus | GGV-L | NC_018482.1 | 471 | 629 |
| Lassa_virus | LASV-S | LT601602.1 | 1545 | 1602 |
| Lujo_virus | LUJV-S | NC_012776.1 | 1373 | 1443 |
| Mopeia_virus | MOPV-S | NC_006575.1 | 1524 | 1646 |
| Mobala_virus | MOBV-S | AY342390.1 | 1561 | 1623 |
| Ippy_virus | IPPYV-S | NC_007905.1 | 1538 | 1595 |
| Junin_virus | JUNV-S | KU978800.1 | 1538 | 1628 |

| | | | | |
|---|---|---|---|---|
| Machupo_virus | MACV-S | NC_005078.1 | 1580 | 1669 |
| Chapare_virus | CHAPV-S | EU260463.1 | 1514 | 1608 |
| Sabiá_virus | SABV-S | U41071.1 | 1526 | 1619 |
| Tacaribe_virus | TACV-S | MT081316.1 | 1521 | 1632 |
| Pichinde_virus | PICV-S | NC_006447.1 | 1564 | 1650 |
| Whitewater_Arroyo_virus | WWAV-S | NC_010700.1 | 1506 | 1576 |
| Amapari_virus | AMAV-S | NC_010247.1 | 1508 | 1606 |
| Cupixi_virus | CPXV-S | AF512832.1 | 1498 | 1592 |
| Pirital_virus | PIRV-S | NC_005894.1 | 1570 | 1659 |
| Paraná_virus | PARV-S | NC_010756.1 | 1577 | 1647 |
| Flexal_virus | FLEV-S | NC_010757.1 | 1583 | 1680 |
| Allpahuayo_virus | ALLV-S | NC_010253.1 | 1719 | 1788 |
| Bear_Canyon_virus | BCNV-S | NC_010256.1 | 1509 | 1590 |
| Latino_virus | LATV-S | NC_010758.1 | 1638 | 1727 |
| Oliveros_virus | OLIV-S | NC_010248.1 | 1674 | 1770 |
| Lymphocytic_choriomeningitis_virus | LCMV-S | NC_077807.1 | 1445 | 1575 |
| Golden_gate_virus | GGV-S | NC_018483.1 | 1445 | 1615 |
| Boa_Arenavirus_NL | BoANL-L | KC508670.1 | 470 | 621 |
| Boa_Arenavirus_NL | BoANL-S | KC508669.1 | 1419 | 1567 |
| Tamiami_Virus | TAMV-L | NC_010702.1 | 368 | 445 |
| Tamiami_Virus | TAMV-S | NC_010701.1 | 1539 | 1719 |
| Guanarito_Virus | GUAV-L | NC_005082.1 | 368 | 455 |
| Guanarito_Virus | GUAV-S | NC_005077.1 | 1514 | 1608 |
| California_Academy_Sciences_Virus | CASV-L | NC_018484.1 | 424 | 515 |
| California_Academy_Sciences_Virus | CASV-S | NC_018481.1 | 1235 | 1422 |
| University_Helsinki_Virus_1 | UHV1-L | KF297881.1 | 472 | 631 |
| University_Helsinki_Virus_1 | UHV1-S | KF297880.1 | 1343 | 1596 |
| Wenzhou_Virus | WENV-L | NC_026019.1 | 335 | 443 |
| Wenzhou_Virus | WENV-S | NC_026018.1 | 1533 | 1594 |

**Supplementary Information 2:** Percentage identity matrix for LCMV and GGV IGR sequences calculated using ClustalW12.1 (Madeira et al., 2024).

| | **LCMV L-IGR** | **LCMV S-IGR** | **GGV L-IGR** | **GGV S-IGR** |
|---|---|---|---|---|
| **LCMV L-IGR** | 100.00 | 56.94 | 51.75 | 54.90 |
| **LCMV S-IGR** | 56.94 | 100.00 | 57.14 | 47.54 |
| **GGV L-IGR** | 51.75 | 57.14 | 100.00 | 53.33 |
| **GGV S-IGR** | 54.90 | 47.54 | 53.33 | 100.00 |

**Supplementary Information 3:** Structure identity percentage calculated using BEAGLE alignment (Mattei et al., 2015) for cryo-OrbiSIMS guided 2D models of LCMV and GGV IGRs. (A) Pairwise comparison of the cryo-OrbiSIMS models of the IGRs and (B) Individual comparison of the cryo-OrbiSIMS models with the *in silico* predicted structure for the respective IGR using Fold algorithm (Bellaousov et al., 2013). The identity percentage is computed as the fraction of paired bases encoded with an identical BEAR character. z-scores (reported in brackets) are the reference statistic measures computed using as background, the distribution of the scores obtained by aligning unrelated RNA sequences (Mattei et al., 2015). Positive z-scores indicate that the 2D structures are more similar than expected, while negative z-scores indicate that they are less similar than expected. A z-score > 3 is considered significant.

(A) Pairwise structure identity percentage matrix for cryo-OrbiSIMS models

| | **LCMV L-IGR** | **LCMV S-IGR** | **GGV L-IGR** | **GGV S-IGR** |
|---|---|---|---|---|
| **LCMV L-IGR** | - | 19.35 (-0.484) | 21.66 (1.007) | 2.3 (-0.244) |
| **LCMV S-IGR** | 19.35 (-0.484) | - | 80.56 (1.201) | 30.56 (0.001) |
| **GGV L-IGR** | 21.66 (1.007) | 80.56 (1.201) | - | 10.18 (1.111) |
| **GGV S-IGR** | 2.3 (-0.244) | 30.56 (0.001) | 10.18 (1.111) | - |

(B) Pairwise structure identity percentage of cryo-OrbiSIMS models (columns) when compared to the respective *in silico* predictions(rows)

| | **LCMV L-IGR** | **LCMV S-IGR** | **GGV L-IGR** | **GGV S-IGR** |
|---|---|---|---|---|
| **LCMV L-IGR** | 51.61 (7.141) | | | |
| **LCMV S-IGR** | | 77.78 (4.557) | | |
| **GGV L-IGR** | | | 75.45 (8.839) | |
| **GGV S-IGR** | | | | 28.89 (4.292) |

**Supplementary Information 4:** Analysis of LCMV L-IGR RNA using optical tweezers unfolding experiments. A total of 60 force extension curves (FECs) were used for this analysis from 10 individual molecules. (A-B) 2D renderings of the (A) *in silico* predicted and (B) cryo-OrbiSIMS-guided models of LCMV L-IGR. Both these structures have high structural identities within the individual stem-loop elements, except in SL1, which is significantly shorter by more than 4 nm in the cryo-OrbiSIMS model compared to *in silico* prediction. (C) Mapping of the individual unfolding events in Figure 2 (main text) to cryo-OrbiSIMS model of LCMV L-IGR. This unambiguously distinguishes the contour length for SL1 to match the cryo-OrbiSIMS model rather than the *in silico* prediction, validating the experimentally derived structure for this IGR.

(A) *In silico* prediction

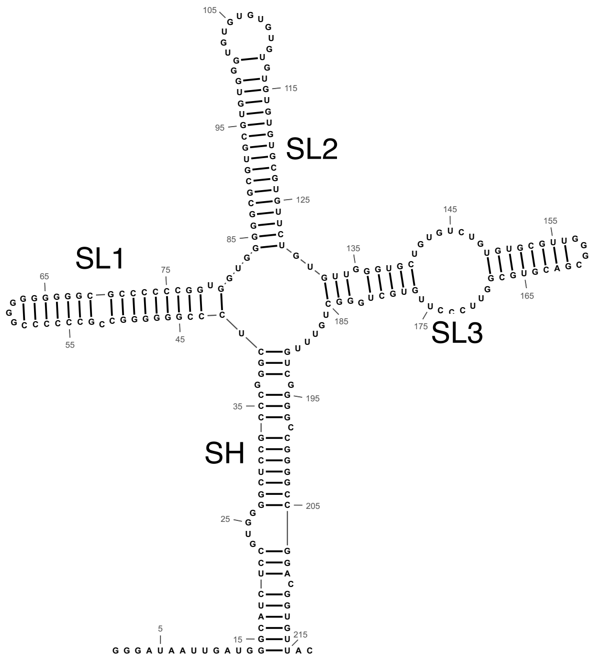


(B) cryo-OrbiSIMS model

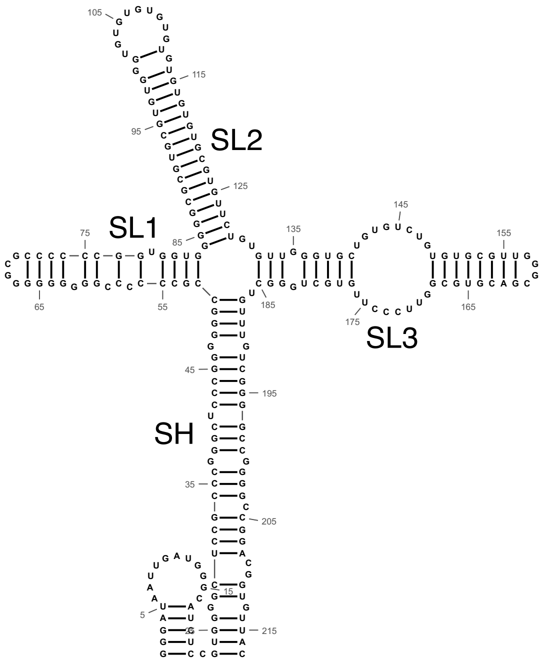


| Element | Bases | nm |
|---|---|---|
| SH | 63 | 37.17 |
| SL1 | 39 | 23.01 |
| SL2 | 46 | 27.14 |
| SL3 | 54 | 31.86 |

| Element | Bases | nm |
|---|---|---|
| SH | 63 | 37.17 |
| SL1 | 32 | 18.88 |
| SL2 | 46 | 27.14 |
| SL3 | 54 | 31.86 |

(C) OT data mapping on LCMV L-IGR Cryo-OrbiSIMS Model

| | **Bases** | **Length (nm)** | **Cumulative Unfolding length (nm)** | **Mapped Unfolding event** |
|---|---|---|---|---|
| Unstructured | 15 | 8.85 | | |
| SH | 61 | | 35.99 | First (SH + R1) |
| B1 | 8 | 4.72 | 4.72 | |
| B2 | 6 | 3.54 | 8.26 | |
| B3 | 17 | 10.03 | 18.29 | |
| B4 | 19 | 11.21 | 29.5 | |
| B5 | 11 | 6.49 | 35.99 | |
| SL1 | 32 | | 18.88 | Last |
| SL2 | 46 | | 27.14 | Second or Third |

| | | | | |
|---|---|---|---|---|
| U1 | 34 | 20.06 | 20.06 | |
| U2 | 12 | 7.08 | 27.14 | |
| SL3 | 54 | | 31.86 | Second or Third (SL3 – R1) |
| R1 | 8 | 4.72 | 4.72 | |
| R2 | 26 | 15.34 | 20.06 | |
| R3 | 20 | 11.8 | 31.86 | |

**Supplementary Information 5:** (a) Ten lowest energy cryo-OrbiSIMS models and (b) *in silico* predictions (Bellaousov et al., 2013) of the GGV S-IGR RNA without the leading IVT linker sequence. (c) BEAGLE comparison of the lowest energy cryo-OrbiSIMS model and *in silico* predicted structure (Ballesio et al., 2025; Mattei et al., 2015). These structures have 20.23% identity (fraction of paired bases encoded with an identical BEAR character) and 52.02% similarity (fraction of paired bases encoded with two different BEAR characters belonging to the same RNA structural element, e.g. two different characters encoding for stems with different lengths), *p-value* 2E-04, indicating that despite some equivalent elements, overall the structures are significantly different from each other. The colour coding highlights sub-substructures that are equivalent between the two models and are not intended to be conservation measures.

> GGV S-IGR Sequence
GCACCACGUGCCCACCCUCUGGUUGUUUGUUUCCCGGAAUGGUAGUCGUGGUUUGU
UUUGUUUGCAGCUGGGUAAGAAAGCAACACCACGUGGCCCACCCUCUGGUUGUUUG
UUUCCCGGUCCGCCCGCCCUGGGUGCUUUGUUCGGUCCGGGCUUGGUUGUUUGGA
CCCGUC

(a) Ten lowest energy Cryo-OrbiSIMS models of GGV S-IGR RNA without IVT linker

>ENERGY = -101.4
((((...))))....................................................(((((((((.(((.(((((((.....(((...)))......))))))).))))))))).((((((...)))
)))...))).((((((((((......))))))).)))..

>ENERGY = -100.0
..((((((............((..........))............))))))...........(((((((((.(((.(((((((.....(((...)))......))))))).))))))))).((((((...)
)))))...))).((((((((((......))))))).)))..

>ENERGY = -98.7
((((...))))....................................................(((((((((.(((.(((((((((...)))..(((.....))))))))).))))))))).((((((...))
))))...))).((((((((((......))))))).)))..

>ENERGY = -97.1
.....(((.................................(((..(............)..))).((((((.(((.(((((((.....(((...)))......))))))).))))))))).((((((...)))
))).........(((((((((......))))))))))))).

>ENERGY = -96.7
((((...)))).........((((((.....................................))))))(((((((.(((((((((((.(((...)))....))).))).))))).......((((((...))
)))).)))))))(((((((((......)))))))))....

>ENERGY = -95.6

(((((..(((........(.((..........)).)........................((.((((((.(((.(((((((.....(((...)))......))))))).)))))))))..)).)))....
))))).......(((((((((......)))))))))....

>ENERGY = -95.6

.....((((((((.......((((((..................................)))))))))).((((..(((((((.(((...)))....))).))))..)))).(((..((((((...))
))))..)))...(((((((((......))))))))))).

>ENERGY = -95.6

(((((.((.(((........)))......................((((...........((.((((((.(((.(((((((.....(((...)))......))))))).)))))))))..))))))..)
)))))).......(((((((((......)))))))))....

>ENERGY = -94.9

(((....(((((((......((........................((((...........((.((((((.(((.(((((((.....(((...)))......))))))).)))))))))..)))))))))
))))))..))).((((((((((......))))))).)))..

>ENERGY = -94.2

((((((((............((..........))............)))))...........))).((((((.(((.(((((((.....(((...)))......))))))).)))))))))..(((((..((
(((((...))))))))..))))).(((((.....)).)))..

(b) In silico predictions of GGV S-IGR RNA without IVT linker

>ENERGY = -56.2

..((((((.((..(((.((((((...........)))))..))).)))))))).(((.......)))((((((.(((.(((((..((((...((....))...))))))))).))))))))).(((((..
...))))).........(((((((((......)))))))))....

>ENERGY = -54.2

((((...)))).(((((...((.((...((...((((...(((...((((((((((((.((((((......)))))))))))).))))))..))).(((....)))..........))))...)).)))
)..))))).........(((((((((......)))))))))....

>ENERGY = -54.0

(((((....((.......((((...........(((((.(((....((((((((((((.((((((......)))))))))))).))))))....)))...)))))...........))))........))...
.))))).......(((((((((......)))))))))....

>ENERGY = -53.2

(((....(((((((.(((((((...........)))))..))....((((((((((((.((((((......)))))))))))).))))))).(((.......((((...........))))........))).
))))))))..))).((((((((((......))))))).)))..

>ENERGY = -52.1

((((...))))(((.....)))...........(((((.(((....((((((((((((.((((((......)))))))))))).))))))....)))...)))))............((((((((((..
(((((.......)))))..)))))..........))))).)..

>ENERGY = -51.2

((((...))))......(((((...........)))))..(((...((((((((((((.(((((......)))))))))))).))))))..)))(((((...((............(((.....))))).
.)))))........(((((((((......)))))))))....

>ENERGY = -51.1

((((...))))(((.....)))...........(((((.(((....((((((((((((.(((((......)))))))))))).))))))....)))...)))))..........(((((.((((((...
..)))..........))).)))))...((((.....))))....

>ENERGY = -50.9

..((((((.((..(((.(((((...........)))))..))).)))))))).(((.......)))........((.((((((.(.(((.(((((..(((....((..........)).)))..).)))))..
))))))))).)).(((((((((......)))))))))....

(c)

Cryo-OrbiSIMS model

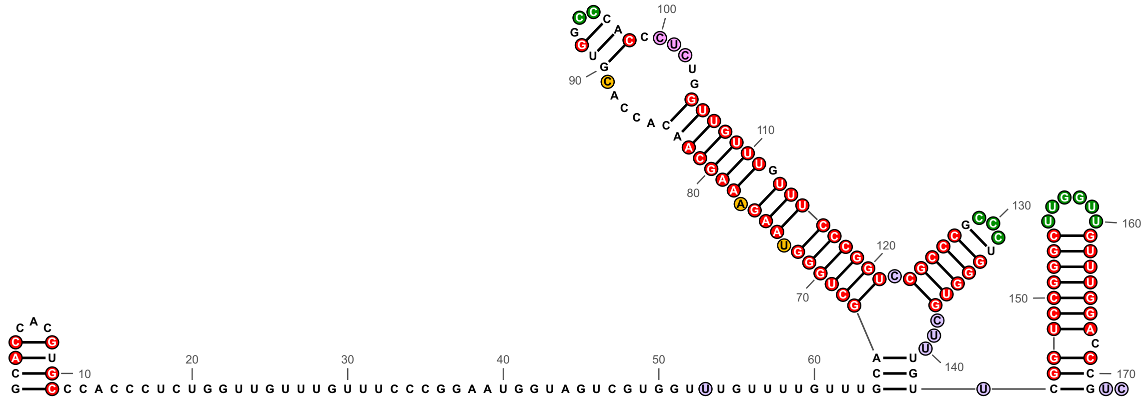

In silico prediction

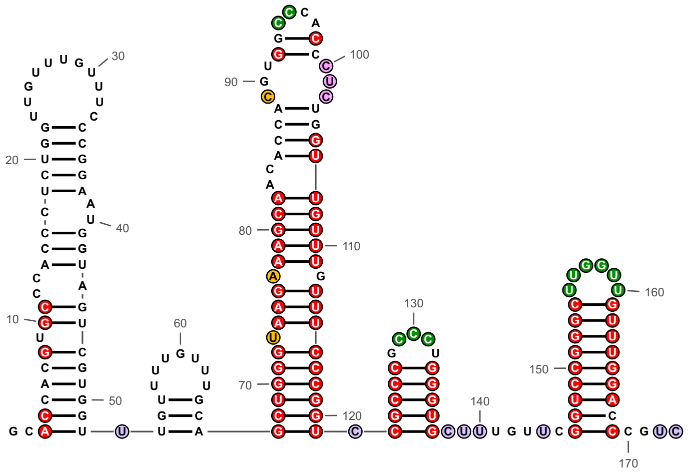

**Supplementary Information 6:** Pairwise Root Mean Squared Deviation (RMSD in Å) calculated using UCSF Chimera (Meng et al., 2006) for cryo-OrbiSIMS guided 3D models of LCMV and GGV IGRs.

| | **LCMV L-IGR** | **LCMV S-IGR** | **GGV L-IGR** | **GGV S-IGR** |
|---|---|---|---|---|
| **LCMV L-IGR** | 0 | 46.83 | 64.65 | 50.24 |
| **LCMV S-IGR** | 46.83 | 0 | 23.26 | 23.73 |
| **GGV L-IGR** | 64.65 | 23.26 | 0 | 60.19 |
| **GGV S-IGR** | 50.24 | 23.73 | 60.19 | 0 |

**Supplementary Information 7:** Graph Representation of RNA structures. Cryo-OrbiSIMS model of LCMV L-IGR is provided here as an example. In this framework, helices (filled boxes) and stem-loops (unfilled boxes) containing them are coded as nodes and the structural features connecting them, such as bulges and loops are coded as edges. In our graph construction, we also heuristically weighted the edges according to whether they would aid coaxial stacking of the two helices they connect. Thus, hairpin loops are weighted 0 as they would cap a helix and not aid in coaxial stack, whereas bulges and internal loops are weighted 1.5 as they are most likely to aid stacking. The edges are also inversely penalised for length, as shorter connectors are likely to lead to structural stability. Helices and stem-loops may have a parent-child relationship. For example, SL1 in LCMV L-IGR is one stem-loop and contains S9-S13 individual helical segments connected by bulges. Centrality metrics, such as strength with k-core, are used to quantify the presence of coaxial stacks in the graph topology. Strength (colour scale) sums up the weights of the edges connected to node and k-core (size of node) measures if the node is embedded in highly interconnected regions. Thus, in densely connected networks such as our RNA graphs, strength and k-core are highly correlated. This means that in the graph representation, central nodes that also likely form coaxial stacks are coloured more magenta and are also larger in size for easy visualisation.

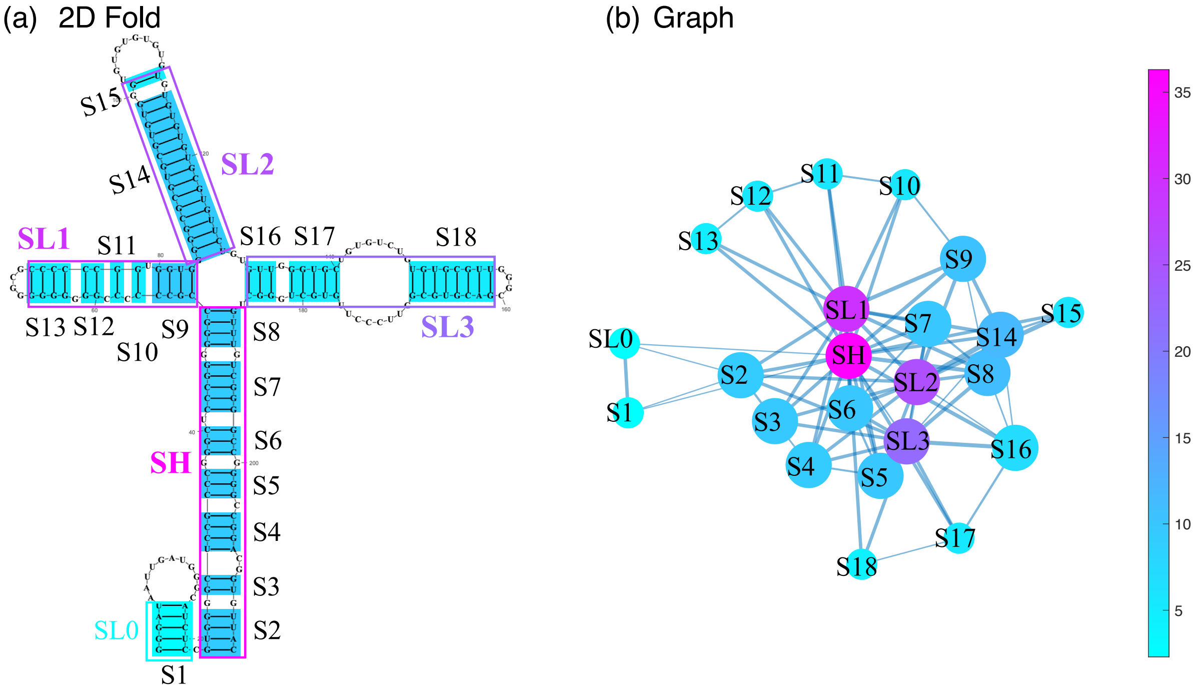

**Supplementary Information 8:** Distribution of graph metrics across the arenavirus IGR space. (a-b) Majority of IGRs (75.8%) exhibit high graph densities (≥ 0.6) and over 70% of the nodes are embedded in k-cores indicated densely connected topologies. (c) List of number of central v/s peripheral stem-loop segments identified across the 58 arenavirus IGRs analysed. One peripheral segment identified in LCMV L-IGR, GGV L-IGR and GGV S-IGR each is due to our spacer sequence *in vitro* template design that has introduced an additional non-native stem-loop at the 5’ end of these IGRs (see main text **Figure 2**).

(a)

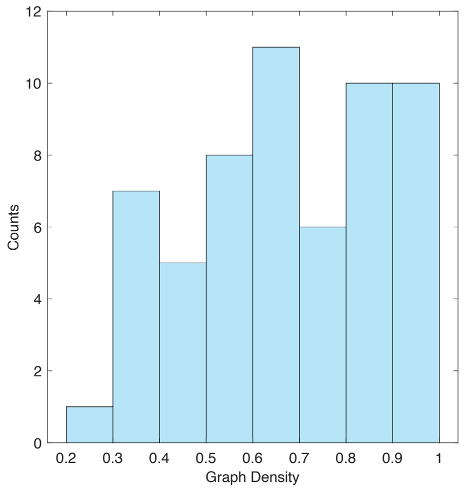


(b)

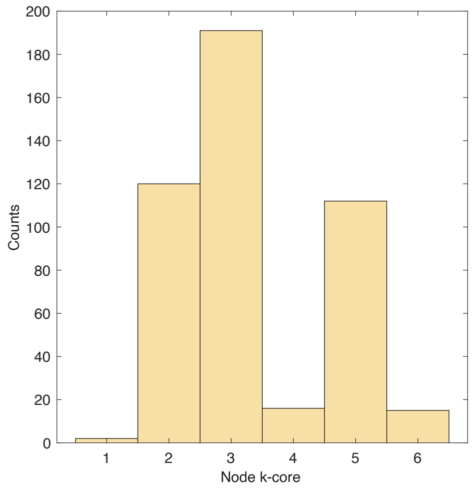


(c)

| ID | IGR | #Total | #Central | #Peripheral |
|---:|---:|---:|---:|---:|
| NC_010249.1 | ALLV-L | 2 | 0 | 0 |
| NC_010253.1 | ALLV-S | 3 | 1 | 0 |
| NC_010251.1 | AMAV-L | 4 | 1 | 0 |
| NC_010247.1 | AMAV-S | 3 | 1 | 2 |
| NC_010255.1 | BCNV-L | 2 | 1 | 1 |
| NC_010256.1 | BCNV-S | 1 | 0 | 0 |
| KC508670.1 | BoANL-L | 5 | 2 | 1 |
| KC508669.1 | BoANL-S | 2 | 2 | 1 |
| NC_018484.1 | CASV-L | 3 | 1 | 2 |
| NC_018481.1 | CASV-S | 2 | 1 | 0 |
| EU260464.1 | CHAPV-L | 3 | 2 | 0 |
| EU260463.1 | CHAPV-S | 3 | 1 | 1 |
| AY216519.2 | CPXV-L | 3 | 3 | 0 |
| AF512832.1 | CPXV-S | 2 | 1 | 0 |

| | | | | |
|---|---|---|---|---|
| NC_010759.1 | FLEV-L | 1 | 1 | 0 |
| NC_010757.1 | FLEV-S | 2 | 1 | 1 |
| NC_018482.1 | GGV-L | 5 | 2 | 2 |
| NC_018483.1 | GGV-S | 6 | 3 | 3 |
| NC_005082.1 | GUAV-L | 1 | 1 | 0 |
| NC_005077.1 | GUAV-S | 2 | 1 | 0 |
| NC_007906.1 | IPPYV-L | 4 | 1 | 1 |
| NC_007905.1 | IPPYV-S | 1 | 1 | 0 |
| KU978799.1 | JUNV-L | 2 | 1 | 1 |
| KU978800.1 | JUNV-S | 3 | 1 | 0 |
| LT601601.1 | LASV-L | 2 | 2 | 1 |
| LT601602.1 | LASV-S | 2 | 0 | 0 |
| NC_010760.1 | LATV-L | 3 | 2 | 1 |
| NC_010758.1 | LATV-S | 2 | 1 | 1 |
| NC_077806.1 | LCMV-L | 5 | 4 | 1 |
| NC_077807.1 | LCMV-S | 2 | 0 | 0 |
| NC_012777.1 | LUJV-L | 4 | 1 | 0 |
| NC_012776.1 | LUJV-S | 1 | 1 | 0 |
| NC_005079.1 | MACV-L | 1 | 1 | 0 |
| NC_005078.1 | MACV-S | 2 | 1 | 0 |
| DQ328876.1 | MOBV-L | 2 | 1 | 1 |
| AY342390.1 | MOBV-S | 2 | 0 | 0 |
| NC_006574.1 | MOPV-L | 3 | 1 | 1 |
| NC_006575.1 | MOPV-S | 4 | 1 | 0 |
| NC_010250.1 | OLIV-L | 2 | 2 | 0 |
| NC_010248.1 | OLIV-S | 3 | 1 | 1 |
| NC_010761.1 | PARV-L | 2 | 1 | 0 |
| NC_010756.1 | PARV-S | 2 | 0 | 0 |
| NC_006439.1 | PICV-L | 2 | 0 | 0 |
| NC_006447.1 | PICV-S | 2 | 1 | 1 |
| NC_005897.1 | PIRV-L | 2 | 1 | 1 |
| NC_005894.1 | PIRV-S | 4 | 1 | 1 |
| AY358026.2 | SABV-L | 1 | 1 | 0 |
| U41071.1 | SABV-S | 3 | 1 | 1 |

| | | | | |
|---|---|---|---|---|
| MT081317.1 | TACV-L | 2 | 1 | 0 |
| MT081316.1 | TACV-S | 3 | 2 | 0 |
| NC_010702.1 | TAMV-L | 2 | 1 | 0 |
| NC_010701.1 | TAMV-S | 5 | 1 | 1 |
| KF297881.1 | UHV1-L | 1 | 1 | 0 |
| KF297880.1 | UHV1-S | 6 | 3 | 1 |
| NC_026019.1 | WENV-L | 2 | 1 | 1 |
| NC_026018.1 | WENV-S | 1 | 0 | 0 |
| NC_010703.1 | WWAV-L | 2 | 1 | 0 |
| NC_010700.1 | WWAV-S | 1 | 1 | 0 |

**Supplementary Information 9:** Robustness of graph-based classification of IGR segments. (a) Stability of node classification: Distribution of the coefficient of variation (CV) of eigenvector centrality calculated for each node across all perturbed networks demonstrates that most nodes exhibit low variability, indicating that their centrality is stable under 20% random perturbations in edge weights. (b-c) Consistency of node classification: For each node originally classified as central (b) or peripheral (c), we tracked its classification across all 100 iterations. (b) Central nodes retained their classification in the majority of perturbations, with 95.5% (64/67) remaining central in ≥90% of iterations. (c) Peripheral nodes were moderately stable, with 63.0% (17/27) retaining their classification in ≥90% of iterations. (d) As an external comparator, Distribution of eigenvector centrality CV calculated for each node across all perturbed networks created for 24 distinct flavivirus 3' UTRs. These are dominated by conserved pseudoknot topologies and not expected to form coaxially stacked helices. The distribution shows that node centrality values were substantially less stable for 3' UTRs, indicating that the robust central helical organisation observed for arenavirus IGRs is not an automatic consequence of applying the framework to structured viral RNAs.

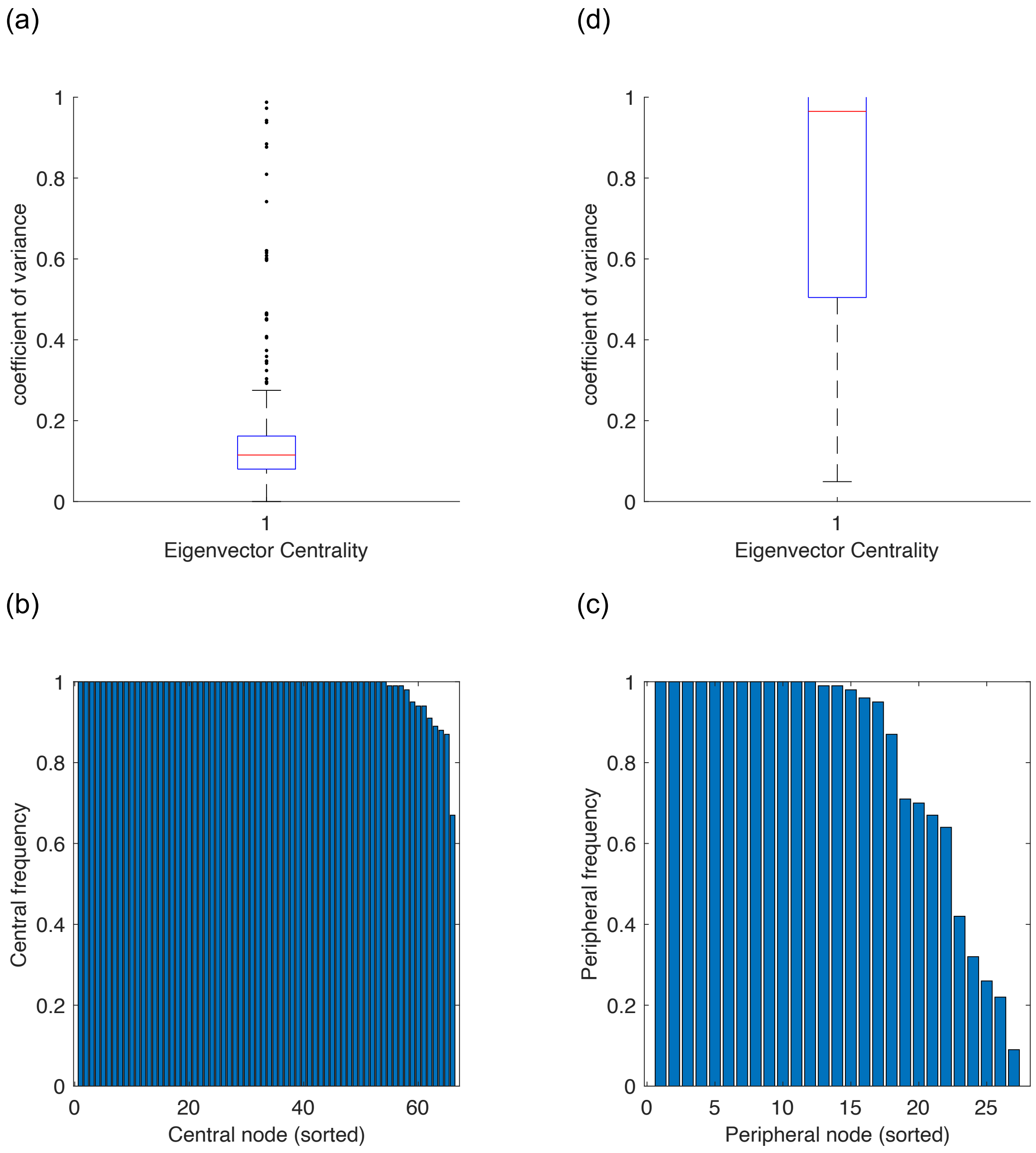

**Supplementary Information 11:** List of significantly enriched proteins with log2(fold-change) > 1.5 at 5% confidence interval in the pull-down interactome of LCMV L-IGR and GGV S-IGR from LCMV infected cell lysate. The enrichment was calculated compared to pull-down experiments with absence of the bait RNA (no IGR negative control).

(A) LCMV L-IGR Interactome

| Accession | log2(fold-change) | -log10(p-value) |
|---:|---:|---:|
| Q08211 | 8.3343 | 4.6622 |
| Q12906 | 7.5179 | 2.1278 |
| P52597 | 5.7754 | 1.8339 |
| O95793 | 5.5138 | 3.2997 |
| P19338 | 5.2837 | 4.8412 |
| Q12905 | 4.8686 | 1.6849 |
| P67809 | 4.8396 | 3.5640 |
| O43390 | 4.5039 | 2.4423 |
| P31943 | 4.4183 | 3.2363 |
| Q9NR30 | 4.3816 | 1.7622 |
| P07910 | 4.3094 | 4.1051 |
| P11940 | 4.2359 | 5.4109 |
| P16401 | 3.7784 | 3.6200 |
| Q8N983 | 3.7663 | 2.4881 |
| Q99729 | 3.7553 | 2.8036 |
| P16402 | 3.6721 | 2.8238 |
| Q9BWV3 | 3.5502 | 2.6085 |
| P61978 | 3.4925 | 2.8323 |
| P38159 | 3.4793 | 2.4883 |
| P16403 | 3.4148 | 2.9738 |
| Q9Y2R9 | 3.2968 | 2.4748 |
| Q9NWU5 | 3.1218 | 1.9138 |
| P52272 | 3.1166 | 4.4259 |
| Q00839 | 3.0977 | 3.5755 |
| P62899 | 2.5473 | 2.4070 |
| P39023 | 2.2048 | 2.8148 |
| Q9BYC9 | 2.0429 | 1.4883 |
| P61247 | 1.8288 | 2.6339 |

| | | |
|---|---|---|
| P50914 | 1.8268 | 2.2541 |
| Q03701 | 1.7769 | 1.3424 |
| P61254 | 1.6271 | 2.7298 |
| P36578 | 1.6032 | 1.8405 |
| Q02878 | 1.5785 | 1.4366 |
| P62851 | 1.5669 | 2.9852 |
| Q92522 | 1.5475 | 2.1346 |

(B) GGV S-IGR Interactome

| Accession | log2(fold-change) | -log10(p-value) | Accession | log2(fold-change) | -log10(p-value) |
|---|---|---|---|---|---|
| Q9NR30 | 6.5900 | 6.7720 | P35268 | 2.2238 | 4.6299 |
| Q9BUJ2 | 6.0977 | 5.3820 | Q5SSJ5 | 2.1089 | 3.2457 |
| P09914 | 5.8280 | 4.7391 | P40939 | 2.1052 | 4.7829 |
| Q08211 | 5.4647 | 9.0714 | Q09666 | 2.0706 | 2.9669 |
| Q12906 | 5.4596 | 8.5289 | P54136 | 2.0070 | 2.0249 |
| P19338 | 4.8340 | 7.3606 | P53396 | 2.0006 | 1.8118 |
| P19525 | 4.8206 | 5.6015 | O43852 | 1.9631 | 2.8965 |
| Q12905 | 4.4990 | 7.9974 | P62899 | 1.9047 | 6.3813 |
| P07910 | 4.1949 | 10.1522 | P61254 | 1.8867 | 8.7322 |
| Q99729 | 3.9646 | 3.5361 | P38159 | 1.8342 | 4.1135 |
| P16401 | 3.9236 | 3.5845 | O95816 | 1.7984 | 3.4373 |
| Q63HN8 | 3.8872 | 6.0520 | P05161 | 1.7964 | 2.2639 |
| P67809 | 3.7374 | 7.3219 | Q15293 | 1.7898 | 4.0306 |
| O43390 | 3.7237 | 4.0464 | P35637 | 1.7626 | 2.9064 |
| O43143 | 3.4710 | 5.8382 | P04406 | 1.7315 | 1.9598 |
| P11387 | 3.4544 | 2.2559 | P31943 | 1.7219 | 3.6350 |
| Q9NZI8 | 3.3877 | 5.9763 | Q1KMD3 | 1.6603 | 4.6423 |
| P26599 | 3.3442 | 4.2736 | P61247 | 1.6389 | 3.8469 |
| Q9GZR7 | 3.1962 | 4.0535 | Q01844 | 1.6226 | 2.2756 |
| Q96HS1 | 3.0761 | 1.9544 | P55265 | 1.6149 | 3.4256 |
| P43243 | 3.0685 | 3.6666 | P84098 | 1.6039 | 4.0744 |
| P22087 | 2.9775 | 3.6706 | P41091 | 1.5432 | 3.9976 |
| Q9NUQ6 | 2.9516 | 3.4749 | P41252 | 1.5318 | 1.6026 |
| P09496 | 2.9413 | 5.9748 | P53007 | 1.5203 | 3.4175 |

| | | |
|---|---|---|
| P42704 | 2.9365 | 7.6370 |
| P52597 | 2.9280 | 3.9832 |
| Q9BQG0 | 2.8991 | 4.1210 |
| Q92522 | 2.8986 | 4.9171 |
| P07305 | 2.8755 | 1.9095 |
| P61978 | 2.8747 | 7.5149 |
| P10412 | 2.6721 | 2.8348 |
| P16402 | 2.6194 | 3.5992 |
| P26368 | 2.5783 | 1.9811 |
| Q52LJ0 | 2.4783 | 3.9250 |
| P16403 | 2.4674 | 2.8028 |
| Q14444 | 2.4660 | 2.5277 |
| P78527 | 2.3187 | 6.7305 |
| Q01130 | 2.3186 | 5.6019 |
| Q00610 | 2.2797 | 5.5799 |
| P29692 | 2.2633 | 2.8529 |
| Q13243 | 2.2425 | 5.2425 |

**Supplementary Information References**


Ballesio, F., Teofani, A., Carrino, C., Catalano, M., Nicolaeasa, M. L., Ausiello, G., Helmer-Citterich, M., & Gherardini, P. F. (2025). BEAGLE 2.0: A Web Server for RNA Secondary Structure Similarity Detection Leveraging SHAPE-directed RNA Structure Determination. *Journal of Molecular Biology*, *437*(15), 169154. https://doi.org/10.1016/j.jmb.2025.169154

Bellaousov, S., Reuter, J. S., Seetin, M. G., & Mathews, D. H. (2013). RNAstructure: Web servers for RNA secondary structure prediction and analysis. *Nucleic Acids Research*, *41*(W1), Article W1. https://doi.org/10.1093/nar/gkt290

Kolberg, L., Raudvere, U., Kuzmin, I., Adler, P., Vilo, J., & Peterson, H. (2023). g:Profiler—Interoperable web service for functional enrichment analysis and gene identifier mapping (2023 update). *Nucleic Acids Research*, *51*(W1), W207–W212. https://doi.org/10.1093/nar/gkad347

Madeira, F., Madhusoodanan, N., Lee, J., Eusebi, A., Niewielska, A., Tivey, A. R. N., Lopez, R., & Butcher, S. (2024). The EMBL-EBI Job Dispatcher sequence analysis tools framework in 2024. *Nucleic Acids Research*, *52*(W1), W521–W525. https://doi.org/10.1093/nar/gkae241

Mattei, E., Pietrosanto, M., Ferrè, F., & Helmer-Citterich, M. (2015). Web-Beagle: A web server for the alignment of RNA secondary structures. *Nucleic Acids Research*, *43*(Web Server issue), W493–W497. https://doi.org/10.1093/nar/gkv489

Meng, E. C., Pettersen, E. F., Couch, G. S., Huang, C. C., & Ferrin, T. E. (2006). Tools for integrated sequence-structure analysis with UCSF Chimera. *BMC Bioinformatics*, *7*(1), 339. https://doi.org/10.1186/1471-2105-7-339

Supek, F., Bošnjak, M., Škunca, N., & Šmuc, T. (2011). REVIGO Summarizes and Visualizes Long Lists of Gene Ontology Terms. *PLoS ONE*, *6*(7), e21800. https://doi.org/10.1371/journal.pone.0021800
